# Molecular early burst associated with the diversification of birds at the K–Pg boundary

**DOI:** 10.1101/2022.10.21.513146

**Authors:** Jacob S. Berv, Sonal Singhal, Daniel J. Field, Nathanael Walker-Hale, Sean W. McHugh, J. Ryan Shipley, Eliot T. Miller, Rebecca T. Kimball, Edward L. Braun, Alex Dornburg, C. Tomomi Parins-Fukuchi, Richard O. Prum, Benjamin M. Winger, Matt Friedman, Stephen A. Smith

## Abstract

Complex patterns of genome and life-history evolution associated with the end-Cretaceous (K– Pg) mass extinction event limit our understanding of the early evolutionary history of crown group birds [1-9]. Here, we assess molecular heterogeneity across living birds using a technique enabling inferred sequence substitution models to transition across the history of a clade [10]. Our approach identifies distinct and contrasting regimes of molecular evolution across exons, introns, untranslated regions, and mitochondrial genomes. Up to fifteen shifts in the mode of avian molecular evolution map to rapidly diversifying clades near the Cretaceous-Palaeogene boundary, demonstrating a burst of genomic disparity early in the evolutionary history of crown birds [11-13]. Using simulation and machine learning techniques, we show that shifts in developmental mode [14] or adult body mass [4] best explain transitions in the mode of nucleotide substitution. These patterns are related, in turn, to macroevolutionary shifts in the allometric scaling relationship between basal metabolic rate and body mass [15, 16]. In agreement with theoretical predictions, this scaling relationship appears to have weakened across the end-Cretaceous transition. Overall, our study provides evidence that the Chicxulub bolide impact [17] triggered integrated patterns of evolution across avian genomes, physiology, and life history that structured the evolutionary potential of modern birds.

## Introduction

It has been over forty years since Alvarez et al. [17] provided chemical evidence indicating that the Cretaceous-Paleogene (K–Pg) mass extinction was associated with an extraterrestrial impact. Subsequent research has confirmed and refined our understanding of this abrupt and cataclysmic event (e.g., [7, 18-21]). Mounting evidence suggests that the K–Pg extinction event triggered convergent patterns of life-history evolution [22]. For example, some lineages experienced a transient “Lilliput effect” in which average body sizes became smaller, likely through faunal sorting, dwarfing, or miniaturization [23-27]. While great effort has been devoted to investigating extinction patterns among various groups across the K–Pg boundary [17, 21, 28], the impact of the end-Cretaceous mass extinction event on the genomes of surviving lineages has received little attention.

Given that life-history parameters such as body mass, generation length, and metabolic rates have been linked to different aspects of molecular evolution [29, 30], convergent patterns of life-history evolution are expected to leave an imprint of the K–Pg extinction event in the genomes of surviving lineages [4, 6]. Only a few studies of animal taxa have attempted to directly investigate how the aftermath of the K-Pg mass extinction event shaped genome evolution (e.g., [4, 8, 31-34]). For example, the evolution of polyploidy may be associated with the K–Pg transition in plants [35, 36], and avian substitution rates may have increased due to extinction-related size-selectivity [4].

Many studies attempting to connect events in Earth’s history to patterns of genome evolution rely on inferences from molecular clock analyses (e.g., [4, 35]). These approaches can reveal heterogeneous patterns in the tempo of molecular evolution (e.g., [6, 37, 38]), but typically assume that the underlying data evolved according to the expectations of a uniform substitution model. If this assumption is violated, time-homogeneous models may obscure important evolutionary patterns. Nevertheless, techniques that enable substitution models to vary across a clade’s evolutionary history have not yet seen widespread adoption in the macroevolution literature (e.g., [39-42]). Detecting model transition points on a phylogeny may provide evidence of evolutionary transitions in the “mode,” or process that generated the observed data [43] – a concept that, when combined with hypothesis testing, has been termed “phylogenetic natural history” [15, 44]. Thus, investigating patterns of model shifts in the context of genome and life–history evolution may reveal unknown links between Earth’s history and evolutionary processes.

Here, we combine approaches from molecular systematics and phylogenetic comparative methods to investigate molecular model heterogeneity across the avian tree of life. We apply a stepwise approach to estimating where molecular substitution model parameters have shifted across a clade’s evolutionary history, implemented in *Janus* [10]. Our analyses reveal well-supported shifts in estimated equilibrium base frequencies across exons, introns, untranslated regions (UTRs), and mitochondrial genomes. Remarkably, model shifts are mostly constrained to previously hypothesized clade originations associated with the K–Pg boundary. This pattern indicates a temporal interval early in crown bird evolutionary history in which the mode of avian genomic sequence evolution rapidly diversified, consistent with an “early burst” macroevolutionary process (e.g., [12, 13, 43, 45, 46]). We examine how this pattern of molecular model shifts is related to macroevolutionary changes in avian life-history variation and focus on aspects of breeding ecology, development, senescence, and metabolism that are thought to have experienced intense selection or relaxation of evolutionary constraints during the K–Pg transition (e.g., [4-6, 47, 48]). Specifically, we investigate the hypothesis that molecular shifts indicate shifts in the adaptive landscape [49] or evolutionary allometries [15]. Our findings demonstrate a roadmap for linking molecular shifts inferred from the comparative analysis of genomic sequences with macroevolutionary patterns detected from the fossil record.

## Results

### Molecular model shifts

We report details of dataset assembly as supplementary material. Fifteen molecular model shifts are identified on total-clades estimated to have originated near the K–Pg boundary [1, 3, 50, 51] (Figure 1, Supplementary Figure 1, 7a-d). Considering overlap across data types, *Janus* detects thirteen phylogenetic regimes (one ancestral + twelve derived) that are required to explain the heterogeneity in modes of sequence evolution across exons, introns, UTRs, and mtDNAs, relative to the [51] MRL3 topology (Figure 1). Most molecular shifts are concordant with the origins of diverse ancient clades previously recognized at ordinal or superordinal taxonomic ranks. These include Notopaleognathae, Tinamiformes, the sister clade to Tinamiformes (in the MRL3 tree; an unnamed clade uniting Rheiformes, Casuariiformes, and Apterygiformes), Neognathae, Columbea, Passerea, Otidae (i.e., Otidimorphae + Strisores, *sensu* Wagler 1830, Jarvis et al. 2014), the sister clade to Otidae (in the MRL3 tree; the remainder of Neoaves), Aequornithes, Coraciimorphae, Psittaciformes, and Passeri (Supplementary Table 1). No false positives are detected in simulations under homogeneous model conditions (Supplementary Appendix).

**Figure 1.**
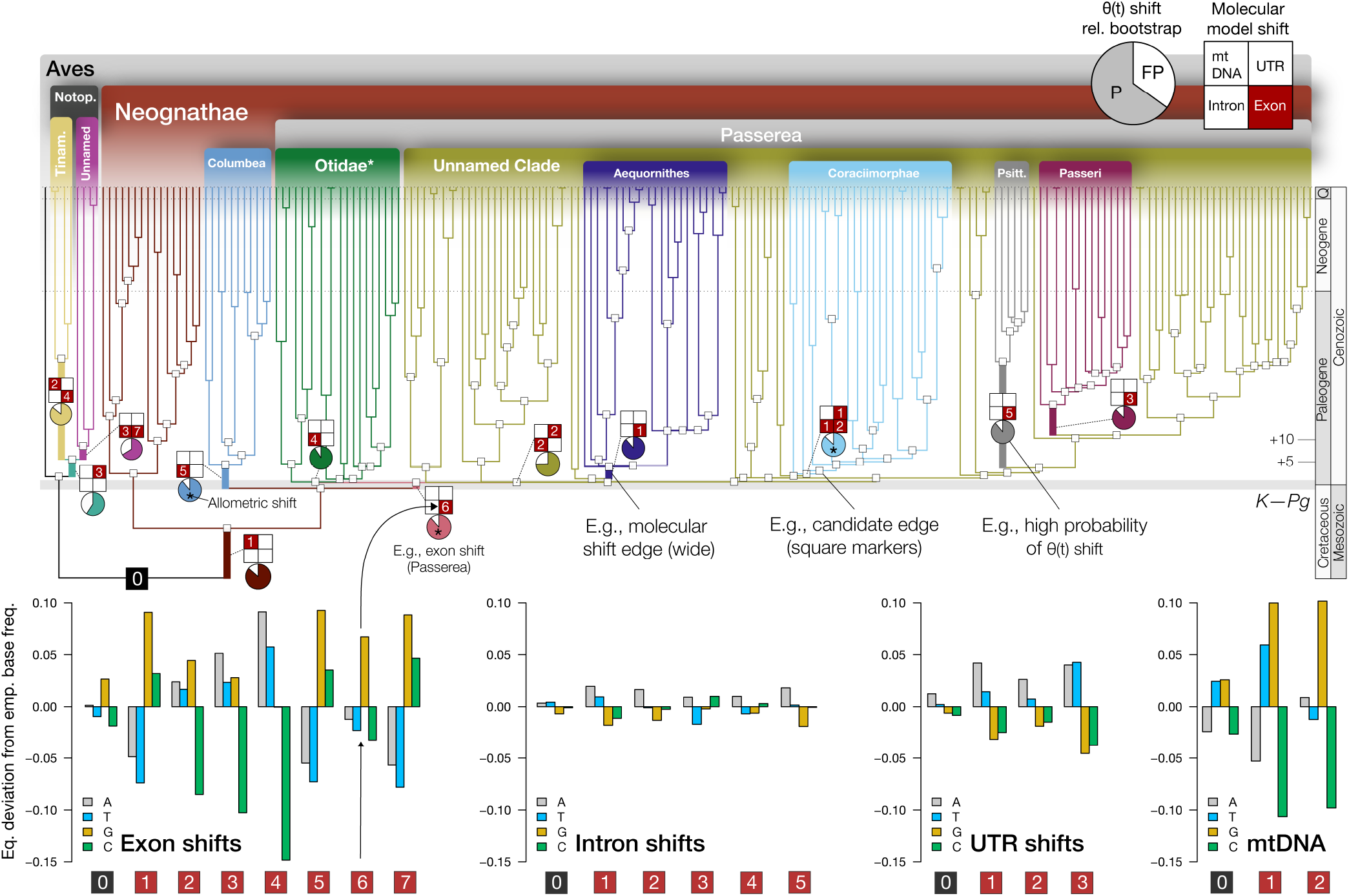
Model shifts across data types. Thirteen phylogenetic regimes encompassing seventeen model shifts are required to explain heterogeneity in equilibrium base frequencies across genetic data types. Fifteen shifts are detected at nodes with stem ages within ∼5 Ma of the K–Pg boundary [51]. Top: the aggregate signal of molecular model shifts across nuclear and mitochondrial data types identified by *Janus*, mapped onto the MRL3 supertree, with the ancestral regime “0” in black (^*^Otidae = Otidimorphae + Strisores, *sensu* Wagler 1830, Jarvis et al. 2014). Patterns of molecular model shifts across data types are summarized as 2×2 grids (Supplementary Figure 3, 7). Numeric labels at each grid position correspond to a particular shift in a specific data type. Pie charts summarize the positive detection rate (“P”) for shifts in trait optima θ(*t*) across eight life-history traits relative to the null false positive (“FP”) rate (e.g., ℓ1ou detection rate / (ℓ1ou detection rate + false positive rate), under AICc; see Supplementary Figure 4). Below: estimated magnitude of shifts in equilibrium base frequency relative to the empirical base frequencies for a given taxon partition for each data type, ordered by dataset size (also see Supplementary Figure 7a-d). Edges with well-supported shifts in metabolic allometry are labeled with an asterisk, with the most substantial support observed for Coraciimorphae (*pp* = 98%, see results and Figure 3).

For every case, our approach identifies molecular shifts with 100% of the model weight when considering a shift’s existence and location, indicating that shifts have a robust statistical signal. Several molecular shifts fall on the GC-AT axis (Figure 1, Supplementary Figure 7a-d) of compositional variation, with the most pronounced shifts occurring in exon data, followed by mtDNA, UTRs, and introns. We detect the highest frequency and magnitude of compositional shifts in our large exon dataset (Figure 1). Nonetheless, there is no trend relating dataset size to the magnitude of inferred substitution parameters (e.g., the relatively small mtDNA data set indicates shifts with significant deviations in estimated equilibrium base frequencies). We observe similar deviations from empirical frequencies among coding (exon, mtDNA) when compared to non-coding sequences (intron, UTR) (Figure 1, see Discussion).

### Life-history evolution

In general, analyses of life-history data support the hypothesis that molecular model shifts coincide with shifts in the evolutionary optima of life-history traits. In this context, optima [θ(*t*)] are equilibrium values within trait space that a lineage evolves towards under stabilizing selection and genetic drift. The detection rate for shifts in θ(*t*) occurring on candidate edges, normalized relative to the false positive rate under mvBM, is consistently high (Figure 1); all candidate cases are supported under AICc or pBIC criteria, but not both in a few cases (e.g., median of 76.2-87.2%, see Supplementary Appendix for additional detail). Analyses with a Random Forest machine learning approach are consistent with those from the model-based analyses reported above (further detail reported in the Supplementary Appendix). Avian developmental mode (“ChickPC1”) [14], followed by adult body mass, are consistently the most important predictors of taxon partitions identified by molecular model shifts (AUC = 0.94). By contrast, traits reflecting substrate or dietary preferences are ranked as relatively unimportant, except for granivory, which is ranked fourth after average clutch size (Figure 2, Supplementary Figure 5, 6).

**Figure 2.**
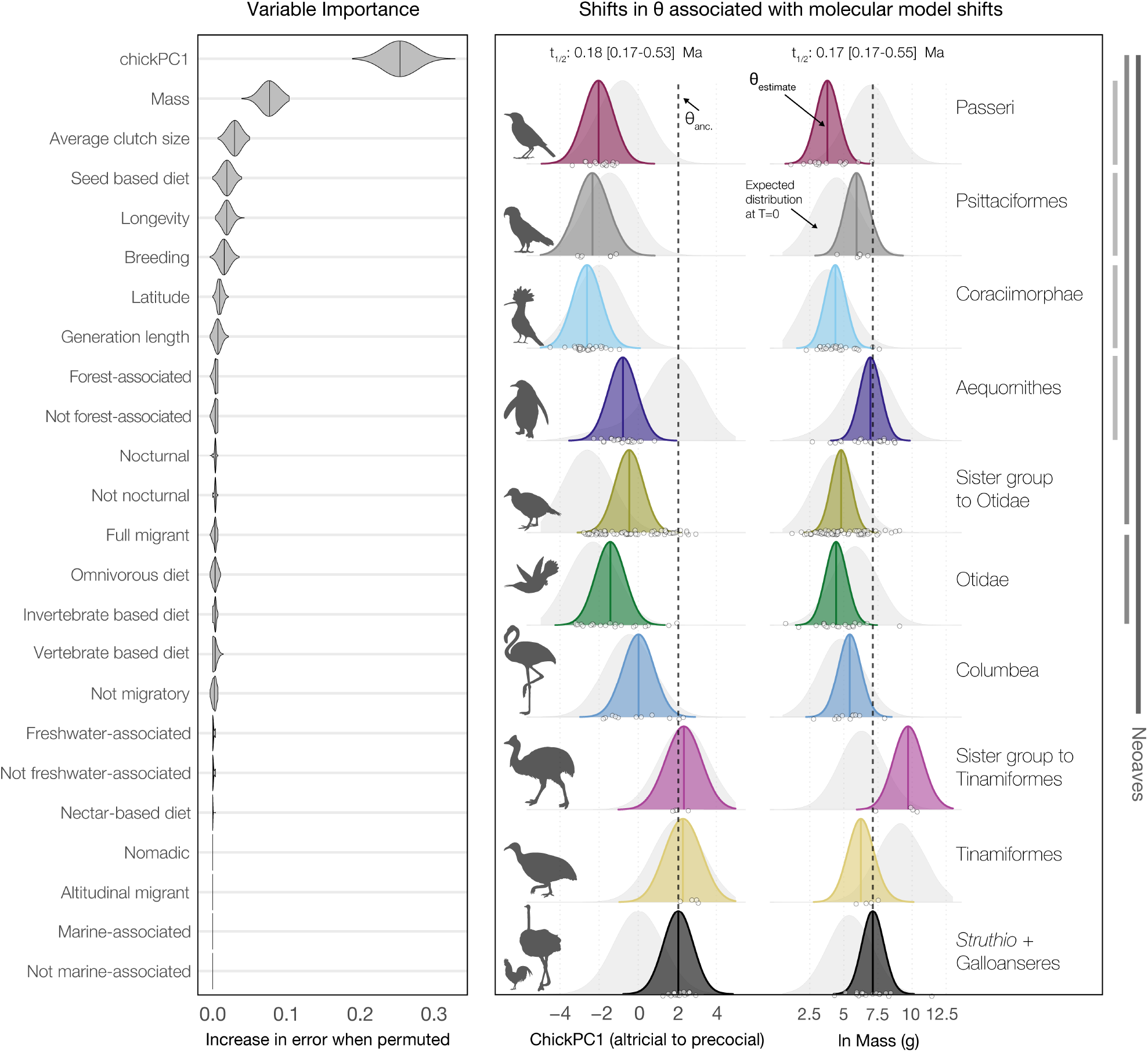
Life-history traits associated with molecular model shifts. Left: permutation-based variable importance for life-history traits reported in [52] and [14]. With a Random Forest classifier, we identified variation in avian developmental mode (“ChickPC1” from [14]) and adult body mass as closely associated with taxon partitions recognized by *Janus* in an analysis of exon data (Supplementary Appendix). Right: estimates of trait optima θ(*t*) when anchored with molecular model shifts from nuclear genetic data (100 parametric bootstraps; colors and labels match Figure 1). Background distributions (light gray) indicate expected values of θ(*t*) at T = 0. For reference, vertical lines mark θ_ancestral_ and phylogenetic levels within Neoaves (e.g., demarking lineages with probable pre- and post-Cretaceous originations). Molecular model shifts are often associated with shifts toward increased altriciality at hatching or decreased adult body mass (also see Supplementary Figures 5, 6).

Within Neoaves, molecular model shifts are broadly associated with θ(*t*) shifts toward increased altriciality at hatching or decreased adult body mass relative to θ_ancestral_ (7/7 and 6/7, respectively, Figure 2). Although with an overall lower θ(*t*) than θ_anc_, Aequornithes and Psittaciformes show derived increases in body mass θ(*t*), along with derived shifts toward increased altriciality θ(*t*) (Figure 2, right). Outside Neoaves, developmental mode θ(*t*) within Palaeognathae is not clearly associated with molecular model shifts. For body mass, however, θ(*t*) for Tinamiformes is similar to θ_anc._, while its unnamed sister clade shows a marked increase in θ(*t*) relative to θ_anc_(Figure 2). Notably, an alternative set of analyses estimating θ(*t*) separately for *Struthio*+root indicates all molecular shifts, including those within Palaeognathae, are associated with derived decreases in body mass or ChickPC1 θ(*t*) (Supplementary Figure 6 and supplemental discussion). These patterns support the hypothesis that molecular model shifts indicate evolutionary shifts in life-history θ(*t*).

### Metabolic allometry

Bayesian models of metabolic allometry indicate that deviations from a prior expectation of 3/4 power law scaling are associated with molecular model shifts close to the K–Pg boundary. Modal estimates for slope (*β*_mass_) range from 0.68 (∼⅔) to 0.84 (∼⅘), and intercept (*β*_0_) from − 4.32 to −3.17 (Figure 3, Supplementary Table 1). These values are similar to those previously estimated for avian and mammalian subclades [53-55]. Compared to life-history traits like mass or developmental mode (Figure 2, Supplementary Figure 5, 6), the evolution of metabolic scaling parameters appears more uncertain, with broad posterior estimates in some cases (e.g., Palaeognathae, Figure 3). Nevertheless, these analyses reveal that the origins of several K–Pg-associated subclades within Neoaves (e.g., Passerea) coincide with a shift toward overall lower body mass, as well as lower slope and higher intercept terms (e.g., under 10 kg as noted in [53, 56]). However, shifts in the allometric optima of individual candidate subclades are not always in a consistent direction (Figure 3).

**Figure 3.**
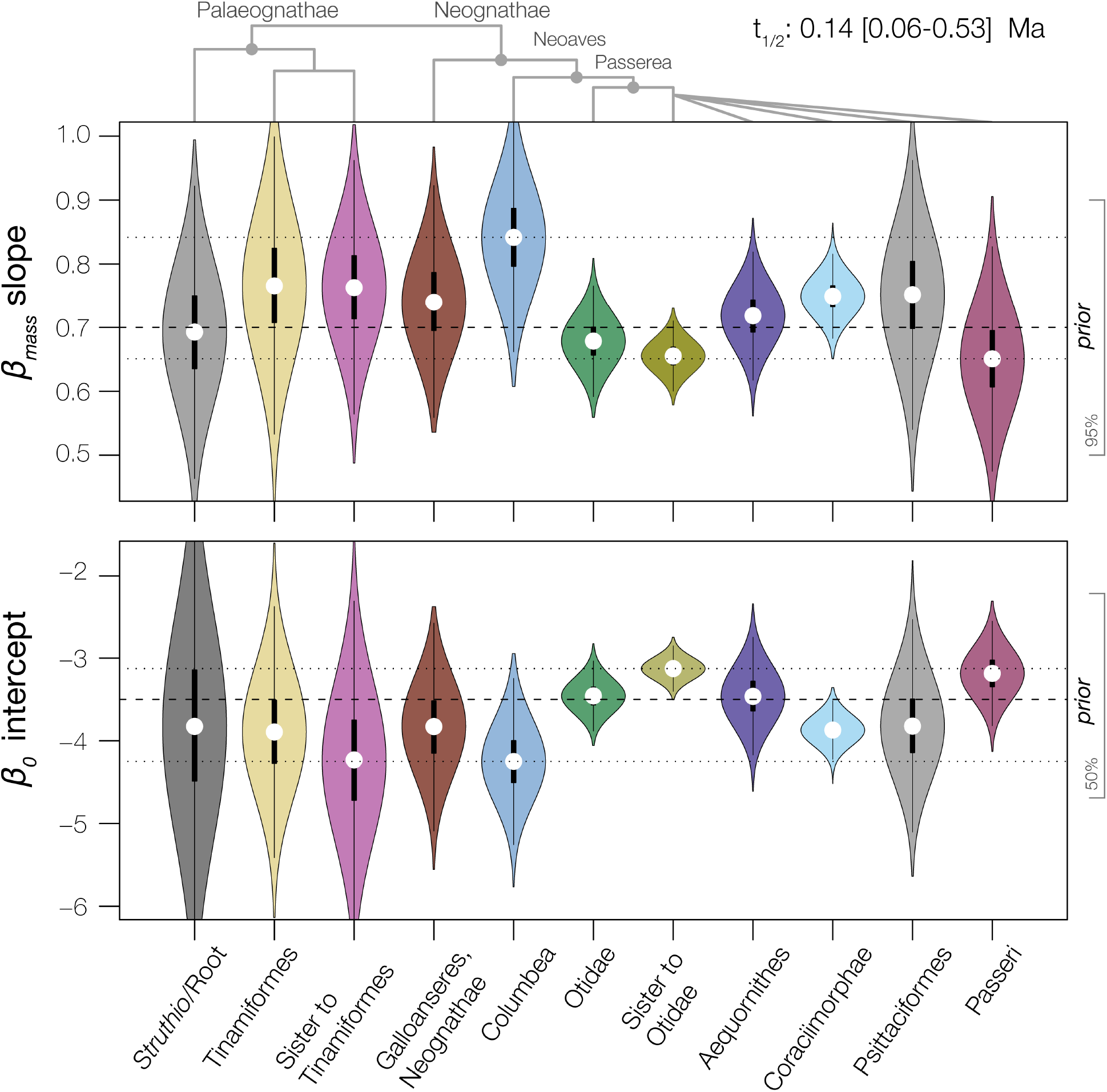
Molecular model shifts are associated with a range of avian metabolic allometries. We analyzed metabolic data under a Bayesian framework to generate posterior estimates of intercept (*β*_0_) and slope (*β*_mass_) coefficients from an evolutionary allometric regression model. Here, we depict the posterior estimates for slope and intercept across a fixed allometric shift model recapitulating molecular model regimes across all data types (single branch internodes reflecting Passerea and Notopalaeognathae not shown; see Supplementary Table 1). Median parameter estimates are indicated with white dots. Horizontal dashed lines indicate the prior mean (prior density intervals are shown on the right vertical axis for reference); dotted lines indicate the range of median parameter estimates. A shift toward lower slope and higher intercept characterizes lineages with probable post-Cretaceous origin (e.g., Passerea).

Seven edges detected at a 10% posterior probability cutoff reflect K–Pg-associated clade originations (Figure 3, Supplementary Table 1). Under a more conservative threshold, only three candidate edges are detected with moderate to strong support, including the unnamed sister clade of Otidae (*pp*∼38%), Columbea (*pp*∼39%), and Coraciimorphae (*pp*∼98%) (Figure 1). The diverse clade Coraciimorphae is the only candidate group for which molecular model shifts are detected across all nuclear genetic data types. Overall, metabolic parameter estimates are consistent with the hypothesis that allometric shifts in avian metabolism may be associated with molecular model shifts.

## Discussion

Unraveling interactions among significant events in Earth’s history and macroevolutionary patterns is a fundamental and persistent problem in evolutionary biology. Our study investigates patterns of sequence evolution and the adaptive radiation of birds near the K–Pg boundary. We demonstrate how proximity to the K–Pg boundary increases the probability of molecular model shifts (Figure 1, Supplementary Figure 1a-d), linking a major mass extinction to the macroevolution of the avian genome. By anchoring a series of phylogenetic comparative models with molecular model shifts, we find evidence that shifts in the mode of genomic evolution were likely concurrent with shifts in the evolutionary optima θ(*t*) of important avian life-history traits (Figure 2), as well as shifts in metabolic allometric slope *β*_mass_ and intercept *β*. Broadly, model shifts in genomic sequences identify shifts toward increased altriciality or smaller adult body mass, consistent with the hypothesis of a K–Pg-associated “Lilliput effect” [4].

Our examination of metabolic allometry provides insight into the biological consequences of size evolution; transitions toward smaller sizes are correlated with weaker scaling relationships between metabolic rate and body mass (e.g., along the Neoaves-Passerea backbone; Figure 3, [4]). This pattern implies that the energetic costs of evolutionary increases in size are reduced in clades with a smaller average body mass. In a scenario like the K–Pg extinction, during which networks of ecological competitors were reset [57], the survivorship of clades with smaller body sizes—and weaker associations between metabolic rate and body mass—may have thus facilitated the evolution of a range of physiological strategies in the early Cenozoic (e.g., [58-60]). This deduction is consistent with theoretical and empirical advances which predict transitions toward harsher environments with increased extrinsic mortality drive the evolution of lower metabolic scaling exponents because of selection to maximize lifetime reproduction [16, 61]. These associated phenomena lead to earlier maturation and faster growth [16], aligning with our inference of increased altriciality associated with the K–Pg extinction (e.g., [14, 62, 63]).

### Recognizing early bursts

Lineages can enter novel adaptive zones during diversification due to ecological, geographic, or phenotypic opportunities [43, 64]. The aftermath of mass extinctions, especially those of short duration, may present all three classes of opportunities, resulting in recovery faunas that experience “early bursts” of lineage and character diversification [45, 65]. If opportunities for diversification become more constrained as ecospace fills, rates of morphological evolution and lineage accumulation should decline, with the fastest rates of change restricted to a short interval following the mass extinction event [11, 12, 66]. Accordingly, we expect initially high rates of evolution to generate outsized disparity early in post-extinction adaptive radiations.

An exclusive focus on rates of change, however, (e.g., [13]) precludes recognition of early burst patterns based on other sources of disparity [67]. Our approach diagnoses a *molecular* early burst of disparate processes of nucleotide sequence evolution. It is, therefore, conceptually more similar to paleontological approaches examining patterns of disparity [46, 67, 68] than it is to comparative techniques estimating rates of change in quantitative or molecular characters [4, 13, 69]. While many studies have quantified early burst phenomena through patterns of morphological evolution or rates of lineage diversification, we show that ancient diversification events may impart a signature of genomic disparity that remains detectable in surviving lineages for tens of millions of years. In this context, the molecular model shifts we identify reveal “genomic fossils” that describe canalized macroevolutionary regimes, within which sequence evolution is aligned along contrasting axes of substitutional variation.

### A novel dimension of avian adaptive radiation

The detection of numerous model shifts within a ∼5 Ma interval of the K–Pg boundary (Figure 1) is likely not a coincidence – and instead reveals a canonical “early burst” pattern in which ecological disparity *and* lineage disparity accumulated rapidly in the early history of crown birds [45, 70]. Given that the uncertainty in estimated molecular divergence times typically exceeds 5 Ma [1, 3, 4, 50, 51, 71], a conservative interpretation of available divergence time estimates does not reject the hypothesis that these events were closely linked (Figure 1, Supplementary Figure 1). We emphasize that *Janus* does not have access to information about the absolute timing of divergence events; it is, therefore, striking that shifts cluster in temporal proximity on a well-justified time-calibrated phylogeny [51] (Figure 1, Supplementary Figure 1).

Consistent with this pattern, models of quantitative trait evolution estimate short phylogenetic half-lives (t_1/2_ = ln(2)/α, e.g., t_1/2_ body mass ∼ 0.18 [0.17-0.52] Ma, Figure 2). Such short intervals imply a scenario with rapid character displacement followed by a relatively stationary process (assuming a median generation length of three years [52] reflects a t_1/2_ of ∼ 60,000 generations). This interpretation is prompted by a conservative view of the fossil record, which currently only supports limited crown bird diversification in the Late Cretaceous (e.g., [4, 6, 50]), and short circum-K—Pg internodes. Our results, therefore, support the hypothesis that developmental and life-history traits were canalized early in crown bird evolutionary history [2, 14, 72], partitioning avian bauplan and higher-taxa (e.g., [67, 73, 74]). Short t_1/2_may also partly explain deviations between estimated equilibrium and empirical base frequencies in these data (Figure 1). Considering the coincidence of molecular model shifts with shifts in the evolutionary optima of life-history traits (e.g., Supplementary Figure 4), both phenomena likely reflect integrated evolutionary responses to the post-Cretaceous adaptive landscape and a permanent shift to new adaptive zones (e.g., [75]).

While many mechanisms link substitution rates to the life-history spectrum [30], we have limited intuition about how the mode of molecular evolution may relate to life-history variation. One idea proposed to explain variation in DNA compositional heterogeneity links a recombination process known as GC-biased gene conversion (gBGC) to generation length, effective population size, and co-varying life-history traits [10, 76-81]. Increases in N_e_ [8] and decreased generation lengths [4] could plausibly explain some of the patterns we observe (e.g., higher N_e_ is predicted to increase the efficacy of gBGC [79]). The fact that we observe many substitution model shifts in exon data is also consistent with the hypothesis that transcriptionally active regions experiencing more recombination may be more subject to gBGC [82, 83]. Lastly, our observation that coding regions have considerably greater deviations between estimated equilibrium and empirical base frequencies could be due to functional constraints, codon usage bias, recombination, or selection (see supplementary analysis of codon usage and Supplementary Figure 7a-d).

Although we do not investigate it directly, our results imply a mechanism to explain the “data type” effect in studies of avian phylogeny [1, 71, 84, 85], in which phylogenetic analyses of coding or non-coding nucleotide data recover conflicting signals of phylogeny. Time-heterogeneous processes of molecular evolution are expected to introduce systematic error in attempts to resolve phylogeny through model-misspecification (e.g., [6, 84, 86]). We note that the data type exhibiting the weakest deviations between empirical and estimated equilibrium base frequencies are non-coding introns (Figure 1). Thus, our results support the conclusions of [71, 84], who favor non-coding data in the inference of avian phylogeny. At the same time, our results show that non-coding data are likely not immune from these issues. Given that numerous model shifts are detected for both coding and non-coding sequences in a region of the avian phylogeny with uncertain topological relationships [71, 87], future progress on avian phylogeny will hinge on our ability to rule out or better accommodate these potential sources of error.

## Conclusion

Although high-throughput sequencing has clarified the history of many vertebrate clades, early evolutionary events within ancient lineages of crown birds—a group comprising over 10,000 extant species—have long been shrouded in mystery. Our study illuminates how the diversification of modern birds in close association with the end-Cretaceous mass extinction was characterized by rapid shifts across several axes of life-history variation. Directional selection on these parameters across the end-Cretaceous mass extinction event, such as selection towards increased altriciality or decreased adult body mass, appears to have shifted the processes of genome evolution. Ultimately, our results reveal a novel dimension of how one of the most significant events in Earth’s history – the Chicxulub bolide impact – structured the evolutionary potential of surviving lineages and gave rise to the spectacular diversity of birds.

## Methods

### Nuclear Sequence Data Collection and Processing

We re-assembled an existing short-read sequence targeting 394 gene regions across 198 bird species and two crocodilian outgroups from *Prum et al*. [3]. These data were initially collected using target-capture of anchored hybrid enrichment (AHE) loci [88], a set of single-copy regions semi-conserved across vertebrates. We analyzed the existing raw sequencing reads with a common pipeline designed to extract phased exons. First, we removed low-quality regions and adaptor sequences using Trimmomatic v0.36 [89] and merged overlapping reads using FLASH v1.2.11 [90]. We assembled reads for each sample using Trinity v2.11 [91]. We then annotated assemblies by comparing assembled contigs to target loci using blat v36×2 [92]. To ensure we annotated orthologs, we retained only contigs with a reciprocal best-hit match to a target locus. To identify intron-exon boundaries, we used exonerate 2.4.0 to compare the nucleotide sequences of annotated loci to the protein sequences for the exons of each locus, based on protein-coding data and annotations from the zebra finch (*Taeniopygia guttata*, genome assembly bTaeGut1_v1). This approach assumes that intron-exon boundaries are conserved across the avian radiation. Occasionally, mapping of the exon sequence to the nucleotide sequence was discontinuous, suggesting the presence of an intervening non-coding region. In such cases, we retained the highest-scoring contiguous stretch of sequence only.

To identify variable sites, we mapped cleaned reads back to annotated contigs using bwa v0.7.17-r1188 [93] and used GATK v4.1.8 to mark duplicates [94]. We called variants on this alignment using GATK HaplotypeCaller and filtered it to only retain variants with coverage >20x and quality >20. Using this high-quality variant set, we recalibrated the base quality scores in the alignment files using GATK. We then called variants and phased them using HaplotypeCaller. Finally, we exported diplotypes and phased haplotypes per intron and exon in a coding region, masking any sites with coverage <2×. Ultimately, we captured 453 exons, 573 introns, and 213 untranslated regions.

Before alignment, we applied a series of sequential filtering steps to remove remaining short or low-quality fragments. We removed (1) leading and trailing N characters from each fragment, and resulting sequences that were zero length (see locus-filtering.R script), (2) fragments with > 40% N characters, (3) fragments that were < 50 bp long, and (4) whole loci that lacked coverage for at least 10% of the taxa in the dataset.

We aligned phased exon sequences with the Multiple Alignment of Coding Sequences (MACSE) ALigning, Filtering, and eXporting pipeline (ALFIX) [95]. MACSE-ALFIX chains together several programs that perform reading frame aware alignment with MACSE and subsequent alignment filtering with HmmCleaner [96] to remove non-homologous sequence fragments. We aligned phased non-coding sequences with Fast Statistical Alignment (FSA) [97]. We calculated alignment statistics using AMAS [98]. We used trimAl [99] to evaluate the effect of 5%-30% alignment column occupancy filtering on alignment length and the loss of parsimony informative sites. We ultimately filtered our non-coding alignments to require a minimum column occupancy of 5% (i.e., 95% of the sequences in an alignment are allowed to contain a gap for a given site). This procedure, which we believe is conservative, increased the signal-to-noise ratio in these data by removing stretches of unaligned nucleotides (characteristic of FSA alignments) while retaining most of the informative data [100] (also see [101]). Unfiltered alignments and the final filtered dataset are provided as Supplementary Data.

### Mitochondrial Sequence Data Collection and Processing

We ran Mitofinder 1.4 [102] to identify the mitochondrial regions from the previously assembled contigs. For reference mitogenomes, we used complete mitogenomes available in GenBank (Supplementary Table 3). When available, we used a reference from the same order (though for Passeriformes, we used different references for oscines [Passeri] and suboscines [Tyranni]); in a few cases, it was necessary to use a reference from a closely related order (Supplementary Table 3). We then extracted the 13 protein-coding genes and 2 rRNAs from the mitofinder output (final_genes.fasta file). In some cases, limited mitochondrial data were recovered (Supplementary Table 3). In those cases, we searched GenBank for the same or a phylogenetically equivalent species that could be substituted. When no suitable alternative was available from GenBank, we also used mitogenomes assembled from the raw data collected as part of other studies ([103-107], and Braun et al. *in prep*). To increase data coverage in five cases, we generated chimeric sequences using available GenBank data from multiple individuals of the same species (Supplementary Table 3).

Once a set of sequences had been assembled, we performed an initial analysis using these data, combined with a larger set of mitogenomes (Kimball *et al*., *in prep*) to ensure sequences were correctly identified (placed phylogenetically with expected relatives) and did not exhibit unusually long branch lengths, which might suggest assembly errors. To do this, we ran an initial alignment using MAFFT 7.294b [108] using default parameters. This alignment was then analyzed in IQ-TREE 2.1.2 ([109]) using the GTR+I+G4 substitution model with 1000 ultrafast bootstrap replicates ([110]). Lastly, we regenerated alignments for the present study, using the same procedure described above for nuclear coding and non-coding data.

### Phylogenetic framework

To avoid issues of circularity related to inferring molecular patterns and phylogenetic topology from the same molecular dataset and to control for stochastic resolutions of Neoaves (e.g., [87]), our focal analyses use the MRL3 supertree [51] as a topological constraint. This topology balances the signal of phylogeny among several recently published avian genomics datasets, and resolves the seven major higher-level clades identified by [84], as well as most intra-ordinal clades that are robustly supported by [3]. It is also in line with a growing number of studies that have suggested that early diversification events within the avian crown group were associated with the K–Pg boundary [1, 3, 50, 51]. As inference of avian phylogeny is an active area of research [71], we explored how patterns of gene-tree discordance (e.g., [1]) may confound inference of molecular model shifts or potential statistical associations with the K–Pg boundary (Supplementary Appendix, Supplementary Figure 1b).

Molecular model shift analysis with *Janus* takes a rooted input phylogram with branch lengths in substitution units (method details reported in Supplementary Appendix, [10]). To generate starting trees for *Janus*, we used our reprocessed datasets and estimated maximum likelihood branch lengths with IQ-TREE v 2.1.1 [109, 111]. For each data type, we applied an optimal partition model selected with the MFP+MERGE approach in IQ-TREE [112, 113], with each locus defined as the unit for partitioning. We estimated molecular branch lengths separately for exons, introns, untranslated regions, and mtDNAs, but kept the topology fixed across datasets.

The *Janus* algorithm fits models that describe patterns of substitutions irrespective of the absolute timing of divergence events (Supplementary Appendix). Therefore, temporal patterns must be evaluated on a reference timeline. To interpret our model-shift results on a time-calibrated phylogeny, we used congruification [114] with treePL [115] to apply the divergence date estimates from the reduced taxon set analysis presented in [51] to the phylogenetic branch length estimates derived from the present study. The well-constrained divergence estimates from [51] are broadly congruent with those reported across several phylogenomic analyses of independent datasets [1, 3, 50]. These estimates reject the hypothesis that many modern avian clades originated in the Cretaceous and centers the diversification of most superordinal variation within ∼ +/− 5 Ma of the K—Pg boundary (Figure 1). Therefore, our interpretations are conditional on this general divergence time scenario, an area of active research [4, 6, 50] (see discussion).

### Fitting time-heterogeneous models with *Janus*

For nuclear genomic data, we considered the signal across three concatenated datasets (exons, introns, and UTRs). For mitochondrial data, we considered three alternative datasets (all data combined, protein-coding genes combined, and rRNAs combined). Our focus on data type mirrors recent developments implicating this axis of genomic variation as a primary source of phylogenetic incongruence [1, 71, 84]. We fit time-heterogenous substitution models to each dataset with *Janus* (Supplementary Appendix, commit 8952e31d, https://git.sr.ht/~hms/janus). By default, *Janus* optimizes a model which enables shifts in equilibrium base frequencies across a provided phylogram. We set each search to accommodate rate heterogeneity across sites according to a discretized gamma distribution (−g) and to assess model weights for the existence (−u) and location (−l) of model shifts. Simulations indicate that this combination of parameter options has high power (e.g., a negligible false-positive rate) to detect the phylogenetic position of molecular model shifts (Supplementary Appendix, [10]). Considering our genetic dataset’s taxonomic sample, we set the minimum clade size to >|= 4 (−m 4). Thus, the set of possible shift configurations reflects 101 internal nodes spanning ∼77 Ma (square markers in Figure 1; e.g., with postorder traversal, any node (excluding the root) with >|= 4 descendant edges).

### Life-history and metabolism datasets

To assess how the configuration of molecular model shifts detected with *Janus* may be related to life-history variation, we considered how life history might vary across multiple dimensions (e.g., [4, 116]). We assembled two life-history datasets to minimize the amount of missing data in each analysis. The first dataset focused on quantitative life-history traits and was compiled from the [52]. These data include body mass, modeled generation length, latitude centroid, mean clutch size, annual adult survival, age at first breeding, maximum longevity, and categorically coded variables for diet, habitat, and diurnality and migratory status. We also included a metric of avian developmental mode (“ChickPC1”) that describes variation in hatchling state along an altricial to precocial spectrum [14]. These data reflect exact species matches relative to those in the re-assembled nuclear genetic dataset.

The second dataset reflects energetic constraints on life-history variation and includes basal metabolic rates (BMR) expressed in watts and associated body masses. Metabolic rates broadly scale as a ∼3/4 power law function of organism mass and reflect rates of energy flow in and through organisms [16, 54, 117-120]. [15] previously considered the hypothesis that allometric scaling parameters relating BMR and body mass have evolved across the vertebrate tree of life. We apply the same general approach to our sample of avian metabolic diversity (below). We first collected available BMR records from the AnAge senescence database Build 14 [121]. For most of the exact species in the present dataset (and most avian species in general), conspecific BMR data have not been measured. Therefore, we conducted an extensive literature search for each avian family in the molecular dataset and filled in many missing entries by identifying phylogenetically equivalent matches (e.g., at the genus level) for which BMR and mass data were available.

Several downstream analyses required complete datasets, so we used two methods to generate unbiased estimates of missing values under a multivariate Brownian motion process (mvBM). In the case of the larger eight-dimensional breeding ecology dataset, we used *Rphylopars* [122] to fit a variance-covariance matrix (VCV) and to estimate values for missing entries. In the case of the two-dimensional metabolic scaling dataset, we used *mvMORPH* [123] to compare the fit of alternative multi-regime, mvBM models based on the model shift points identified by *Janus*. In the latter case, the values of the imputed data were virtually identical across alternative models (e.g., R^2^ > 0.98), so we selected the model with the lowest AIC score to use for downstream analyses.

### Analysis of life-history data

Using multiple approaches, we investigated the degree to which patterns of life-history variation reflect distinct evolutionary regimes that coincide with molecular model shifts. Several methods have been developed to automatically generate evolutionary hypotheses by identifying an optimized configuration of evolutionary models describing variation in the process of trait evolution (e.g., [124, 125]), but few are expressly multivariate (e.g., [49, 126]). We investigated model heterogeneity across our high-dimensional life-history dataset with the bootstrapping approach implemented in the software ℓ1ou [49]. ℓ1ou uses a phylogenetic lasso method to identify points on a phylogeny where a trait’s optimum value θ(*t*) has shifted, assuming α (the “pull” toward the optimum or adaptation rate) and σ^2^ (the Brownian diffusion rate parameter) are fixed across the tree. The ℓ1ou approach is extended to multiple traits by assuming that traits shift their optimum simultaneously and in the same location on the tree [49].

Conveniently, ℓ1ou allows the researcher to specify a set of candidate edges for the lasso approach to consider for shifts in θ(*t*). This attribute allows us to articulate the specific models we want to compare. We ran ℓ1ou with a constrained set of candidate edges reflecting the 12 candidate shift edges identified across analyses of different molecular data types (Figure 1, Supplementary Table 1). Thus, for a given ℓ1ou analysis of this type, ℓ1ou can infer 0-12 shifts in θ(*t*). We repeated this procedure with the AICc and pBIC information criteria, as recommended by [49], and used 100 bootstrap replicates to assess the positive detection rate for each candidate edge. We emphasize that our intention is not to identify every case where a life-history shift may have occurred across avian phylogeny; our goal is to assess how much statistical support exists for shifts in life-history trait optima that coincide with shifts identified in our analysis of molecular data.

We validated our results through comparison to a null distribution of shift detections reflecting the false positive rate under multivariate Brownian motion (e.g., without shifts in θ(*t*). Using the eight-dimensional variance-covariance matrix (VCV) estimated by *RPhylopars* [122], we simulated 500 null datasets using the function simRatematrix in the R package *ratematrix* [127]. We then analyzed each simulated dataset with ℓ1ou as previously specified. For each candidate edge, we used Fisher’s exact test [128] to assess whether the frequency of positive shift detections in the empirical dataset was significantly greater (one-tailed p-value = 0.05) than the null false-positive rate observed across simulated datasets. Tables of p-values and odds ratios are reported as supplementary material (Supplementary Figure 4, Supplementary Table 2).

Lastly, we investigated which life-history traits are most closely associated with molecular model shifts using a machine learning approach implemented in the *tidymodels* framework [129] (Supplementary Appendix). To explore these results, we fit fixed shift OUM (shifting θ(*t*), with fixed α and σ2; equivalent to that used by ℓ1ou) models using *OUwie* [130]. We used 100 parametric bootstrap replicates to estimate model parameter uncertainty. We also simulated 1000 datasets under each fitted OUM model to visualize the expected distribution of trait values at T (time) = 0 (Figure 2). These models, therefore, reflect the expected shifts in trait optima which coincide with bipartitions identified in analyses of molecular model shifts.

### Analysis of metabolic rate data

To assess whether molecular model shifts are associated with shifts in patterns of metabolic scaling, we assessed support for coincident shifts in metabolic allometry. We utilized the Bayesian phylogenetic framework implemented in the R package *bayou* 2.0 [15, 124]. *bayou* applies a reversible jump Markov Chain Monte Carlo approach to detect the magnitude, number, and phylogenetic position of model shifts. Using *bayou*, we implemented an allometric regression model which relates BMR and body mass logarithmically, and for which slope *β*_mass_ and intercept *β*_0_ evolve under a multi-regime Ornstein Uhlenbeck (OU) process. Here, model shifts reflect shifts in the optimum of the evolutionary allometry between BMR and body mass.

Using the rjMCMC approach in *bayou*, we estimated the posterior probability of an allometric shift occurring along the 12 candidate edges identified by *Janus*. Under a Poisson prior, we specified the mean number of shifts across the phylogeny reflecting 2% of the total edges in the tree (λ = 8) with equal probability. In this context, maximal posterior probability indicates an increase over the prior probability by ∼50%. We ran each analysis across three replicate chains for 10 million iterations, sampling every 1000 iterations. Given that we did not have consistent estimates of measurement error, we followed the approach of [15] and explored alternative analyses assuming a Standard Error of 0.1 or 0.01 for BMR and recovered a negligible impact (not shown).

Our priors for α, σ^2^, *β* _*>mass*_ *>, β*_0_ reflect half-Cauchy or Gaussian expectations:

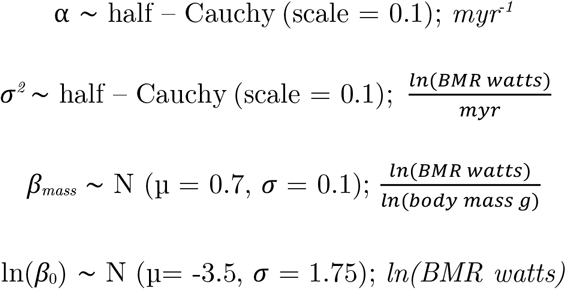

Replicate analyses with rjMCMC identified numerous shifts in the slope and intercept at a posterior probability cutoff of 0.1, after discarding the first 40% of samples as burn-in. Following rjMCMC runs, we re-estimated model parameters on a fixed configuration model reflecting all molecular shifts identified with *Janus* (Figure 3). We assessed model convergence by examining Gelman and Rubin’s R statistic [131] and effective sample sizes across chains and parameters.

## Data Availability

All data or software code generated for the present manuscript is available at the corresponding author’s GitHub repository https://github.com/jakeberv/avian_molecular_shifts and archived at Zenodo pending peer review. *Janus* has been implemented in Golang and C and is available at https://git.sr.ht/~hms/janus and https://git.sr.ht/~hms/hringhorni.

## Acknowledgments

J. S. B. was supported by the University of Michigan Life Sciences Fellows program and the Jean Wright Cohn Endowment Fund at the University of Michigan Museum of Zoology. D. J. F. was supported by UKRI (MR/S032177/1). N. W. H. was funded by the Woolf Fisher Cambridge Scholarship. R. T. K. and E. L. B. were supported by the United States National Science Foundation (DEB 1655683). S. A. S. was supported by the United States National Science Foundation (DBI 1930030 and DEB 1938969). We thank the numerous researchers and museums whose field collection efforts made this research possible. We thank X anonymous reviewers for their feedback and comments. This work was partly supported by the facilities and staff of the University of Michigan Department of Ecology and Evolution Biology and the Yale University Faculty of Arts and Sciences High-Performance Computing Center. We thank the facilitators of the 2017 Evolutionary Quantitative Genetics workshop, including Joe Felsenstein, Brian O’Meara, and Josef Uyeda, for helpful discussions. We thank members of the Winger, Friedman, and Smith labs, Abby Kimmitt, Teresa Pegan, Liam Revell, William Gearty, Jacob Socolar, Jonathan Hughes, and Rafael Marcondes, and Austin Patton for their valuable comments. We thank Mike Braun for providing access to several mitochondrial genomes. Jacob S. Berv wrote the manuscript with input from all co-authors. For the purpose of open access, the authors have applied a Creative Commons Attribution (CC BY) license to any Author Accepted manuscript version arising.

## CRediT (Contributor Roles Taxonomy)

**Jacob S. Berv:** Conceptualization, Methodology, Software, Validation, Formal analysis, Investigation, Resources, Data Curation, Writing - Original Draft, Writing - Review & Editing, Visualization, Supervision, Project Administration, Funding acquisition. **Sonal Singhal:** Conceptualization, Methodology, Software, Validation, Formal analysis, Data Curation, Writing - Review & Editing. **Daniel J. Field:** Conceptualization, Resources, Writing - Review & Editing. **Nathanael Walker-Hale:** Conceptualization, Methodology, Software, Writing - Review & Editing. **Sean McHugh:** Methodology, Formal analysis, Writing - Review & Editing. **J. Ryan Shipley:** Conceptualization, Validation, Investigation, Data Curation, Writing - Review & Editing. **Eliot T. Miller**: Conceptualization, Methodology, Software, Writing - Review & Editing. **Rebecca T. Kimball:** Conceptualization, Formal analysis, Investigation, Resources, Data Curation, Writing - Review & Editing. **Edward L. Braun:** Conceptualization, Resources, Data Curation, Writing - Review & Editing. **Alex Dornburg:** Conceptualization, Methodology, Software, Writing - Review & Editing. **C. Tomomi Parins-Fukuchi:** Conceptualization, Methodology, Software, Writing - Review & Editing. **Richard O. Prum:** Conceptualization, Resources, Funding acquisition, Writing - Review & Editing. **Benjamin M. Winger:** Funding acquisition, Writing - Review & Editing. **Matt Friedman:** Conceptualization, Resources, Funding acquisition, Writing - Review & Editing. **Stephen A. Smith:** Conceptualization, Methodology, Software, Resources, Supervision, Funding acquisition, Writing - Review & Editing.

## Literature Cited

1. Jarvis, E.D., et al., Whole-genome analyses resolve early branches in the tree of life of modern birds. Science, 2014. 346(6215): p. 1320–1331.

2. Ksepka, D.T., et al., Tempo and Pattern of Avian Brain Size Evolution. Current Biology, 2020. 30(11): p. 2026–2036.e3.

3. Prum, R.O., et al., A comprehensive phylogeny of birds (Aves) using targeted next-generation DNA sequencing. Nature, 2015. 526(7574): p. 569–573.

4. Berv, J.S. and D.J. Field, Genomic Signature of an Avian Lilliput Effect across the K-Pg Extinction. Systematic Biology, 2018. 67(1): p. 1–13.

5. Field, D.J., et al., Early Evolution of Modern Birds Structured by Global Forest Collapse at the End-Cretaceous Mass Extinction. Current Biology, 2018. 28(11): p. 1825–1831.e2.

6. Field, D.J., et al., Chapter 5: Timing the Extant Avian Radiation: The Rise of Modern Birds, and the Importance of Modeling Molecular Rate Variation, in Pennaraptoran Theropod Dinosaurs Past Progress and New Frontiers, M. Pittman and X. Xu, Editors. 2020, Bulletin of the American Museum of Natural History: New York. p. 159–181.

7. Longrich, N.R., T. Tokaryk, and D.J. Field, Mass extinction of birds at the Cretaceous– Paleogene (K–Pg) boundary. Proceedings of the National Academy of Sciences, 2011. 108(37): p. 15253–15257.

8. Houde, P., E.L. Braun, and L. Zhou, Deep-Time Demographic Inference Suggests Ecological Release as Driver of Neoavian Adaptive Radiation. Diversity, 2020. 12(4): p. 164.

9. Parins-Fukuchi, C., G.W. Stull, and S.A. Smith, Phylogenomic conflict coincides with rapid morphological innovation. Proceedings of the National Academy of Sciences, 2021. 118(19): p. e2023058118.

10. Smith, S.A., N. Walker-Hale, and C. Parins-Fukuchi, Compositional shifts associated with major evolutionary transitions in plants. bioRxiv, 2022: p. 2022.06.13.495913.

11. Gavrilets, S. and J.B. Losos, Adaptive Radiation: Contrasting Theory with Data. Science, 2009. 323(5915): p. 732–737.

12. Ingram, T., L.J. Harmon, and J.B. Shurin, When should we expect early bursts of trait evolution in comparative data? Predictions from an evolutionary food web model. Journal of Evolutionary Biology, 2012. 25(9): p. 1902–1910.

13. 1. Harmon, L.J., et al., Early bursts of body size and shape evolution are rare in comparative data. Evolution, 2010. 64(8): p. 2385–2396.

14. Ducatez, S. and D.J. Field, Disentangling the avian altricial-precocial spectrum: Quantitative assessment of developmental mode, phylogenetic signal, and dimensionality. Evolution, 2021. 75(11): p. 2717–2735.

15. Uyeda, J.C., et al., The Evolution of Energetic Scaling across the Vertebrate Tree of Life. The American Naturalist, 2017. 190(2): p. 185–199.

16. White, C.R., et al., Metabolic scaling is the product of life-history optimization. Science, 2022. 377(6608): p. 834–839.

17. Alvarez, L.W., et al., Extraterrestrial Cause for the Cretaceous-Tertiary Extinction. Science, 1980. 208(4448): p. 1095–1108.

18. Brugger, J., G. Feulner, and S. Petri, Baby, it’s cold outside: Climate model simulations of the effects of the asteroid impact at the end of the Cretaceous. Geophysical Research Letters, 2017. 44(1): p. 419–427.

19. Chiarenza, A.A., et al., Asteroid impact, not volcanism, caused the end-Cretaceous dinosaur extinction. Proceedings of the National Academy of Sciences, 2020. 117(29): p. 17084–17093.

20. Brusatte, S.L., et al., The extinction of the dinosaurs. Biological Reviews, 2015. 90(2): p. 628–642.

21. Longrich, N.R., B.-A.S. Bhullar, and J.A. Gauthier, Mass extinction of lizards and snakes at the Cretaceous–Paleogene boundary. Proceedings of the National Academy of Sciences, 2012. 109(52): p. 21396–21401.

22. McKinney, M.L., Extinction Vulnerability and Selectivity: Combining Ecological and Paleontological Views. Annual Review of Ecology and Systematics, 1997. 28(1): p. 495–516.

23. Harries, P.J. and P.O. Knorr, What does the ‘Lilliput Effect’ mean? Palaeogeography, Palaeoclimatology, Palaeoecology, 2009. 284(1): p. 4–10.

24. Keller, G. and S. Abramovich, Lilliput effect in late Maastrichtian planktic foraminifera: Response to environmental stress. Palaeogeography, Palaeoclimatology, Palaeoecology, 2009. 284(1): p. 47–62.

25. Lyson, T.R., et al., Exceptional continental record of biotic recovery after the Cretaceous–Paleogene mass extinction. Science, 2019. 366(6468): p. 977–983.

26. Urbanek, A., Biotic crises in the history of Upper Silurian graptoloids: A Palaeobiological model. Historical Biology, 1993. 7(1): p. 29–50.

27. Friedman, M., Ecomorphological selectivity among marine teleost fishes during the end-Cretaceous extinction. Proceedings of the National Academy of Sciences, 2009. 106(13): p. 5218–5223.

28. Wiest, L.A., et al., Terrestrial evidence for the Lilliput effect across the Cretaceous-Paleogene (K-Pg) boundary. Palaeogeography, Palaeoclimatology, Palaeoecology, 2018. 491: p. 161–169.

29. Bromham, L., Why do species vary in their rate of molecular evolution? Biology Letters, 2009. 5(3): p. 401–404.

30. Bromham, L., Causes of Variation in the Rate of Molecular Evolution, in The Molecular Evolutionary Clock. 2020, Springer. p. 45–64.

31. Wu, J., T. Yonezawa, and H. Kishino, Rates of Molecular Evolution Suggest Natural History of Life History Traits and a Post-K-Pg Nocturnal Bottleneck of Placentals. Current Biology, 2017. 27(19): p. 3025–3033.e5.

32. Phillips, M.J., Geomolecular Dating and the Origin of Placental Mammals. Systematic Biology, 2015.

33. Koenen, E.J.M., et al., The Origin of the Legumes is a Complex Paleopolyploid Phylogenomic Tangle Closely Associated with the Cretaceous–Paleogene (K–Pg) Mass Extinction Event. Systematic Biology, 2020. 70(3): p. 508–526.

34. Brown, J.W., et al., Strong mitochondrial DNA support for a Cretaceous origin of modern avian lineages. BMC Biology, 2008. 6(1): p. 1–18.

35. Vanneste, K., et al., Analysis of 41 plant genomes supports a wave of successful genome duplications in association with the Cretaceous–Paleogene boundary. Genome Research, 2014. 24(8): p. 1334–1347.

36. Fawcett, J.A., S. Maere, and Y. Van de Peer, Plants with double genomes might have had a better chance to survive the Cretaceous–Tertiary extinction event. Proceedings of the National Academy of Sciences, 2009. 106(14): p. 5737–5742.

37. Smith, S.A. and M.J. Donoghue, Rates of Molecular Evolution Are Linked to Life History in Flowering Plants. Science, 2008. 322(5898): p. 86–89.

38. Dornburg, A., et al., Relaxed Clocks and Inferences of Heterogeneous Patterns of Nucleotide Substitution and Divergence Time Estimates across Whales and Dolphins (Mammalia: Cetacea). Molecular Biology and Evolution, 2012. 29(2): p. 721–736.

39. Foster, P.G., Modeling Compositional Heterogeneity. Systematic Biology, 2004. 53(3): p. 485–495.

40. Ritchie, A.M., T.L. Stark, and D.A. Liberles, Inferring the number and position of changes in selective regime in a non-equilibrium mutation-selection framework. BMC Ecology and Evolution, 2021. 21(1): p. 39.

41. Romiguier, J., et al., Fast and robust characterization of time-heterogeneous sequence evolutionary processes using substitution mapping. PLoS One, 2012. 7.

42. Dutheil, J.Y., et al., Efficient Selection of Branch-Specific Models of Sequence Evolution. Molecular Biology and Evolution, 2012. 29(7): p. 1861–1874.

43. Simpson, G.G., Tempo and mode in evolution. 1944: Columbia University Press.

44. Uyeda, J.C., R. Zenil-Ferguson, and M.W. Pennell, Rethinking phylogenetic comparative methods. Systematic Biology, 2018. 67(6): p. 1091–1109.

45. Givnish, T.J., Adaptive radiation versus ‘radiation’ and ‘explosive diversification’: why conceptual distinctions are fundamental to understanding evolution. New Phytologist, 2015. 207(2): p. 297–303.

46. Hughes, M., S. Gerber, and M.A. Wills, Clades reach highest morphological disparity early in their evolution. Proceedings of the National Academy of Sciences, 2013. 110(34): p. 13875–13879.

47. Maor, R., et al., Temporal niche expansion in mammals from a nocturnal ancestor after dinosaur extinction. Nature Ecology & Evolution, 2017. 1(12): p. 1889–1895.

48. Hughes, J.J., et al., Ecological selectivity and the evolution of mammalian substrate preference across the K–Pg boundary. Ecology and Evolution, 2021. 11(21): p. 14540–14554.

49. Khabbazian, M., et al., Fast and accurate detection of evolutionary shifts in Ornstein– Uhlenbeck models. Methods in Ecology and Evolution, 2016. 7(7): p. 811–824.

50. Claramunt, S. and J. Cracraft, A new time tree reveals Earth history’s imprint on the evolution of modern birds. Science Advances, 2015. 1(11).

51. Kimball, R.T., et al., A Phylogenomic Supertree of Birds. Diversity, 2019. 11(7): p. 109.

52. Bird, J.P., et al., Generation lengths of the world’s birds and their implications for extinction risk. Conservation Biology, 2020. 34(5): p. 1252–1261.

53. Dodds, P.S., D.H. Rothman, and J.S. Weitz, Re-examination of the “3/4-law” of Metabolism. Journal of Theoretical Biology, 2001. 209(1): p. 9–27.

54. Ballesteros, F.J., et al., On the thermodynamic origin of metabolic scaling. Scientific Reports, 2018. 8(1): p. 1448.

55. Annette E. Sieg, et al., Mammalian Metabolic Allometry: Do Intraspecific Variation, Phylogeny, and Regression Models Matter? The American Naturalist, 2009. 174(5): p. 720–733.

56. Heusner, A.A., Size and power in mammals. Journal of Experimental Biology, 1991. 160(1): p. 25–54.

57. Proche ş Ş., G. Polgar, and D.J. Marshall, K-Pg events facilitated lineage transitions between terrestrial and aquatic ecosystems. Biology Letters, 2014. 10(6): p. 20140010.

58. Jønsson, K.A., et al., Ecological and evolutionary determinants for the adaptive radiation of the Madagascan vangas. Proceedings of the National Academy of Sciences, 2012. 109(17): p. 6620–6625.

59. Schweizer, M., S.T. Hertwig, and O. Seehausen, Diversity versus disparity and the role of ecological opportunity in a continental bird radiation. Journal of Biogeography, 2014. 41(7): p. 1301–1312.

60. Harmon, L.J., et al., Tempo and Mode of Evolutionary Radiation in Iguanian Lizards. Science, 2003. 301(5635): p. 961–964.

61. Glazier, D.S., et al., Ecological effects on metabolic scaling: amphipod responses to fish predators in freshwater springs. Ecological Monographs, 2011. 81(4): p. 599–618.

62. Ricklefs, R.E., R.E. Shea, and I.-H. Choi, INVERSE RELATIONSHIP BETWEEN FUNCTIONAL MATURITY AND EXPONENTIAL GROWTH RATE OF AVIAN SKELETAL MUSCLE: A CONSTRAINT ON EVOLUTIONARY RESPONSE. Evolution, 1994. 48(4): p. 1080–1088.

63. Ricklefs, R.E., ADAPTATION, CONSTRAINT, AND COMPROMISE IN AVIAN POSTNATAL DEVELOPMENT. Biological Reviews, 1979. 54(3): p. 269–290.

64. Schluter, D., The ecology of adaptive radiation. 2000: OUP Oxford.

65. Gould, S.J., The structure of evolutionary theory. 2002: Harvard University Press.

66. Martin, C.H. and E.J. Richards, The Paradox Behind the Pattern of Rapid Adaptive Radiation: How Can the Speciation Process Sustain Itself Through an Early Burst? Annual Review of Ecology, Evolution, and Systematics, 2019. 50(1): p. 569–593.

67. Wagner, P.J., Early bursts of disparity and the reorganization of character integration. Proceedings of the Royal Society B: Biological Sciences, 2018. 285(1891): p. 20181604.

68. Gould, S.J., Wonderful life: the Burgess Shale and the nature of history. 1990: WW Norton & Company.

69. Blomberg, S.P., T. Garland, and A.R. Ives, TESTING FOR PHYLOGENETIC SIGNAL IN COMPARATIVE DATA: BEHAVIORAL TRAITS ARE MORE LABILE. Evolution, 2003. 57(4): p. 717–745, 29.

70. Dornburg, A., et al., THE INFLUENCE OF AN INNOVATIVE LOCOMOTOR STRATEGY ON THE PHENOTYPIC DIVERSIFICATION OF TRIGGERFISH (FAMILY: BALISTIDAE). Evolution, 2011. 65(7): p. 1912–1926.

71. Braun, E.L. and R.T. Kimball, Data Types and the Phylogeny of Neoaves. Birds, 2021. 2(1).

72. Cooney, C.R., et al., Ecology and allometry predict the evolution of avian developmental durations. Nature Communications, 2020. 11(1): p. 2383.

73. Stearns, S.C., The Influence of Size and Phylogeny on Patterns of Covariation among Life-History Traits in the Mammals. Oikos, 1983. 41(2): p. 173–187.

74. Parins-Fukuchi, C., Mosaic evolution, preadaptation, and the evolution of evolvability in apes. Evolution, 2020. 74(2): p. 297–310.

75. Uyeda, J.C., et al., The million-year wait for macroevolutionary bursts. Proceedings of the National Academy of Sciences, 2011. 108(38): p. 15908–15913.

76. Nabholz, B., et al., Dynamic evolution of base composition: causes and consequences in avian phylogenomics. Mol Biol Evol, 2011. 28.

77. Birdsell, J.A., Integrating Genomics, Bioinformatics, and Classical Genetics to Study the Effects of Recombination on Genome Evolution. Molecular Biology and Evolution, 2002. 19(7): p. 1181–1197.

78. Romiguier, J., et al., Contrasting GC-content dynamics across 33 mammalian genomes: Relationship with life-history traits and chromosome sizes. Genome Research, 2010. 20(8): p. 1001–1009.

79. Weber, C.C., et al., Evidence for GC-biased gene conversion as a driver of between-lineage differences in avian base composition. Genome Biology, 2014. 15(12): p. 549.

80. Galtier, N., et al., GC-content evolution in mammalian genomes: the biased gene conversion hypothesis. Genetics, 2001. 159.

81. Bolívar, P., et al., GC-biased gene conversion conceals the prediction of the nearly neutral theory in avian genomes. Genome Biology, 2019. 20(1): p. 5.

82. Aguilera, A., The connection between transcription and genomic instability. The EMBO Journal, 2002. 21(3): p. 195–201.

83. Singhal, S., et al., Stable recombination hotspots in birds. Science, 2015. 350(6263): p. 928–932.

84. Reddy, S., et al., Why Do Phylogenomic Data Sets Yield Conflicting Trees? Data Type Influences the Avian Tree of Life more than Taxon Sampling. Systematic Biology, 2017. 66(5): p. 857–879.

85. Kuhl, H., et al., An Unbiased Molecular Approach Using 3′-UTRs Resolves the Avian Family-Level Tree of Life. Molecular Biology and Evolution, 2020. 38(1): p. 108–127.

86. Jermiin, L.S., et al., The Biasing Effect of Compositional Heterogeneity on Phylogenetic Estimates May be Underestimated. Systematic Biology, 2004. 53(4): p. 638–643.

87. Suh, A., The phylogenomic forest of bird trees contains a hard polytomy at the root of Neoaves. Zoologica Scripta, 2016. 45(S1): p. 50–62.

88. Lemmon, A.R., S.A. Emme, and E.M. Lemmon, Anchored Hybrid Enrichment for Massively High-Throughput Phylogenomics. Systematic Biology, 2012. 61(5): p. 727–744.

89. Bolger, A.M., M. Lohse, and B. Usadel, Trimmomatic: a flexible trimmer for Illumina sequence data. Bioinformatics, 2014. 30(15): p. 2114–2120.

90. Magoč, T. and S.L. Salzberg, FLASH: fast length adjustment of short reads to improve genome assemblies. Bioinformatics, 2011. 27(21): p. 2957–2963.

91. Grabherr, M.G., et al., Full-length transcriptome assembly from RNA-Seq data without a reference genome. Nature Biotechnology, 2011. 29(7): p. 644–652.

92. Kent, W.J., BLAT—The BLAST-Like Alignment Tool. Genome Research, 2002. 12(4): p. 656–664.

93. Li, H., Aligning sequence reads, clone sequences and assembly contigs with BWA-MEM. arXiv preprint arXiv:1303.3997, 2013.

94. Van der Auwera, G.A. and B.D. O’Connor, Genomics in the Cloud: Using Docker, GATK, and WDL in Terra. 2020: O’Reilly Media, Incorporated.

95. Ranwez, V., N. Chantret, and F. Delsuc, Aligning Protein-Coding Nucleotide Sequences with MACSE, in Multiple Sequence Alignment: Methods and Protocols, K. Katoh, Editor. 2021, Springer US: New York, NY. p. 51–70.

96. Di Franco, A., et al., Evaluating the usefulness of alignment filtering methods to reduce the impact of errors on evolutionary inferences. BMC Evolutionary Biology, 2019. 19(1): p. 21.

97. Bradley, R.K., et al., Fast Statistical Alignment. PLOS Computational Biology, 2009. 5(5): p. e1000392.

98. Borowiec, M.L., AMAS: a fast tool for alignment manipulation and computing of summary statistics. PeerJ, 2016. 4: p. e1660.

99. Capella-Gutiérrez, S., J.M. Silla-Martínez, and T. Gabaldón, trimAl: a tool for automated alignment trimming in large-scale phylogenetic analyses. Bioinformatics, 2009. 25(15): p. 1972–1973.

100. Talavera, G. and J. Castresana, Improvement of Phylogenies after Removing Divergent and Ambiguously Aligned Blocks from Protein Sequence Alignments. Systematic Biology, 2007. 56(4): p. 564–577.

101. Tan, G., et al., Current Methods for Automated Filtering of Multiple Sequence Alignments Frequently Worsen Single-Gene Phylogenetic Inference. Systematic Biology, 2015. 64(5): p. 778–791.

102. Allio, R., et al., MitoFinder: Efficient automated large-scale extraction of mitogenomic data in target enrichment phylogenomics. Molecular Ecology Resources, 2020. 20(4): p. 892–905.

103. McCullough, J.M., et al., A Laurasian origin for a pantropical bird radiation is supported by genomic and fossil data (Aves: Coraciiformes). Proceedings of the Royal Society B: Biological Sciences, 2019. 286(1910): p. 20190122.

104. Ericson, P.G.P., et al., Parallel Evolution of Bower-Building Behavior in Two Groups of Bowerbirds Suggested by Phylogenomics. Systematic Biology, 2020. 69(5): p. 820–829.

105. Kimball, R.T., et al., When good mitochondria go bad: Cyto-nuclear discordance in landfowl (Aves: Galliformes). Gene, 2021. 801: p. 145841.

106. Harvey, M.G., et al., The evolution of a tropical biodiversity hotspot. Science, 2020. 370(6522): p. 1343–1348.

107. Smith, B.T., et al., Phylogenomic analysis of the parrots of the world distinguishes artifactual from biological sources of gene tree discordance. Systematic Biology, 2022.

108. Standley, D.M. and K. Katoh, MAFFT Multiple Sequence Alignment Software Version 7: Improvements in Performance and Usability. Molecular Biology and Evolution, 2013. 30(4): p. 772–780.

109. Minh, B.Q., et al., IQ-TREE 2: New Models and Efficient Methods for Phylogenetic Inference in the Genomic Era. Molecular Biology and Evolution, 2020. 37(5): p. 1530–1534.

110. Hoang, D.T., et al., UFBoot2: Improving the Ultrafast Bootstrap Approximation. Molecular Biology and Evolution, 2018. 35(2): p. 518–522.

111. Nguyen, L.-T., et al., IQ-TREE: A Fast and Effective Stochastic Algorithm for Estimating Maximum-Likelihood Phylogenies. Molecular Biology and Evolution, 2014. 32(1): p. 268–274.

112. Kalyaanamoorthy, S., et al., ModelFinder: fast model selection for accurate phylogenetic estimates. Nature Methods, 2017. 14: p. 587.

113. Chernomor, O., B.Q. Minh, and A. von Haeseler, Terrace Aware Data Structure for Phylogenomic Inference from Supermatrices. Systematic Biology, 2016. 65(6): p. 997–1008.

114. Eastman, J.M., L.J. Harmon, and D.C. Tank, Congruification: support for time scaling large phylogenetic trees. Methods in Ecology and Evolution, 2013. 4(7): p. 688–691.

115. Smith, S.A. and B.C. O’Meara, treePL: divergence time estimation using penalized likelihood for large phylogenies. Bioinformatics, 2012. 28(20): p. 2689–2690.

116. Martin, T.E., Avian Life-History Evolution has an Eminent Past: Does it Have a Bright Future? The Auk, 2004. 121(2): p. 289–301.

117. West, G.B., J.H. Brown, and B.J. Enquist. A General Model for the Origin of Allometric Scaling Laws in Biology. in Science. 1997.

118. Brown, J.H., et al., TOWARD A METABOLIC THEORY OF ECOLOGY. Ecology, 2004. 85(7): p. 1771–1789.

119. West, G.B., J.H. Brown, and B.J. Enquist, A General Model for the Origin of Allometric Scaling Laws in Biology. Science, 1997. 276(5309): p. 122–126.

120. Hulbert, A.J., A Sceptics View: “Kleiber’s Law” or the “3/4 Rule” is neither a Law nor a Rule but Rather an Empirical Approximation. Systems, 2014. 2(2): p. 186–202.

121. Tacutu, R., et al., Human Ageing Genomic Resources: Integrated databases and tools for the biology and genetics of ageing. Nucleic Acids Research, 2013. 41(D1): p. D1027–D1033.

122. Goolsby, E.W., J. Bruggeman, and C. Ané, Rphylopars: fast multivariate phylogenetic comparative methods for missing data and within-species variation. Methods in Ecology and Evolution, 2017. 8(1): p. 22–27.

123. Clavel, J., G. Escarguel, and G. Merceron, mvmorph: an r package for fitting multivariate evolutionary models to morphometric data. Methods in Ecology and Evolution, 2015. 6(11): p. 1311–1319.

124. Uyeda, J.C. and L.J. Harmon, A Novel Bayesian Method for Inferring and Interpreting the Dynamics of Adaptive Landscapes from Phylogenetic Comparative Data. Systematic Biology, 2014. 63(6): p. 902–918.

125. Mitov, V., K. Bartoszek, and T. Stadler, Automatic generation of evolutionary hypotheses using mixed Gaussian phylogenetic models. Proceedings of the National Academy of Sciences, 2019. 116(34): p. 16921–16926.

126. Bastide, P., et al., Inference of Adaptive Shifts for Multivariate Correlated Traits. Systematic Biology, 2018. 67(4): p. 662–680.

127. Caetano, D.S. and L.J. Harmon, ratematrix: An R package for studying evolutionary integration among several traits on phylogenetic trees. Methods in Ecology and Evolution, 2017. 8(12): p. 1920–1927.

128. Fisher, R.A., Statistical methods for research workers, 5th ed. Statistical methods for research workers, 5th ed. 1934, Oliver and Boyd: Edinburgh.

129. Kuhn, M. and H. Wickham, Tidymodels: a collection of packages for modeling and machine learning using tidyverse principles. 2020.

130. Beaulieu, J.M., et al., MODELING STABILIZING SELECTION: EXPANDING THE ORNSTEIN–UHLENBECK MODEL OF ADAPTIVE EVOLUTION. Evolution, 2012. 66(8): p. 2369–2383.

131. Gelman, A. and D.B. Rubin, Inference from iterative simulation using multiple sequences. Statistical science, 1992: p. 457–472.

