## Supplementary Appendix for "Molecular early burst associated with the diversification of birds at the K–Pg boundary"

### Molecular and life-history datasets

Our final combined nuclear genetic dataset is a matrix of 400 rows (two alleles per taxon for each locus) and 682,178 aligned base pairs. These data reflect an effective increase in called bases by a factor of ~3.45 and an increase in alignment length by a factor of 1.73 compared with the original dataset from [1], accompanied by increases in the percentage of undetermined and gap characters of less than 10% overall. Only ~14% of the characters in the combined dataset are gaps "-", and ~8% are undetermined characters. The final combined mitochondrial dataset encompasses all 15 targeted regions (13 proteins and 2 rRNAs) and 200 taxa with an average alignment length of 13,769 base pairs (see Supplementary Appendix and Supplementary Table 3). Both life-history datasets reflect the phylogenetic sample of avian lineages in [1]. For the first dataset, we assembled data for body mass (0% missing), modeled generation length (0% missing), latitude centroid (0% missing), mean clutch size (~13.6% missing), annual adult survival (77% missing), age at first breeding (58% missing), maximum known longevity (54.5% missing), and developmental mode ("ChickPC1" – 0% missing). Categorical variables have no missing data (Figure 2, left). For the second dataset highlighting avian metabolism, our procedure to identify phylogenetically equivalent swaps generated ~70% coverage for BMR and 100% for body mass (Supplementary Table 4).

#### Notes on the reassembled molecular dataset

The percentage of undetermined 'N' characters was relatively low in exons (~1.3%) but substantially higher in introns (~24%) and UTRs (~18%). Our final filtered dataset comprised 384 exons (average ~1,198bp), 365 introns (average ~404bp), 147 UTRs (average ~453bp), and 14 loci which were noncoding but did not clearly map to a non-coding genic region (average ~578bp). The reprocessed dataset also reflects an increase in the number of parsimony informative sites by a factor of 2.18 for the phased alignment (~1.8 for a single allele) relative to the [1] dataset. These encouraging results imply that many previously generated phylogenomics datasets may be effectively reprocessed with updated approaches to generate significantly larger and well-annotated resources for studying phylogeny and molecular evolution. Due to computational bottlenecks, our molecular model shift analyses were conducted on sub-sampled sequence datasets representing one sampled allele from each species (we provide the re-assembled datasets as supplementary data). Lastly, the final combined mitochondrial dataset encompassed all 15 targeted regions (13 proteins and 2 rRNAs) and covered 200 taxa with an alignment length of 13,769 base pairs. These data also have a low percentage of missing or undetermined characters (12.8%).

#### Shift detection with *Janus* and model performance

*Janus* has been implemented in both Golang and C, and the source code is available at https://git.sr.ht/~hms/janus and https://git.sr.ht/~hms/hringhorni. The algorithm to detect shifts in stationary frequencies follows a stepwise procedure similar to [2, 3] and requires a rooted tree and matching alignment as input: 1) Estimate a maximum likelihood root composition. 2) Traverse the tree in postorder fashion (i.e., from the tips to the root) and estimate maximum likelihood compositions for subtrees with a minimum number of tips specified by the user. 3) Taking the subtree composition and the root composition for the remainder of the tree, estimate a likelihood and then a Bayesian Information Criterion (BIC) score [4]. 4) Order these compositional shift models for every eligible subtree by BIC. 5) Initiate the final model configuration with root model. Then in a greedy manner, add a shift to the root model based on the previously ordered subtrees, estimate a new BIC, and add the sub-model to the set of models if the new BIC is lower. 6) Discard a sub-model if the updated BIC score is increased by an arbitrary cutoff. Then, using two approaches, we assess the relative support for shifts both with respect to a shift's existence and its location. First, we assess shift existence (-u flag) by evaluating BIC weights for alternative models which include or exclude each proposed shift. Next, we assess the location of proposed shifts (-l flag) by estimating the BIC weights of a proposed shift at the focal node and its two daughters.

Other work showcases a range of simulated conditions under which we evaluate *Janus's* performance; we direct readers to that publication for additional detail [5]. We generally observe that *Janus* is conservative, with negligible false-positive rates after removing poorly supported shifts under the BIC. When configured to estimate uncertainty, *Janus* is also robust to phylogenetic inference error [5].

Here, we assess an additional series of simulated datasets to evaluate the false positive rate for detecting molecular model shifts. Specifically, the molecular phylogeny of Aves is characterized by an overall rapid and early pattern of speciation, in which many super-ordinal clades simultaneously experience early bursts of lineage accumulation [1, 6, 7]. Our simulations are thus designed to assess how an extinction-driven bottleneck, followed by rapid cladogenesis, might impact the inference of molecular model shifts with *Janus*.

First, we take as phylogenetic frameworks the same trees we used as starting trees for *Janus* – these are inferences based on exon, intron, UTR, and mtDNA data sets, as described in the main text. These four trees vary substantially in relative branch lengths and total tree length but are constrained to the same underlying topology. We simulated 100, 2,000bp sequence alignments for each phylogenetic framework under a homogeneous HKY+GC model in IQ-TREE 2.2.0 [8] using AliSim [9]. The relative transition/transversion rate was set to 2:1, and equilibrium base frequencies were randomly assigned for each simulated alignment. The shape parameter for the continuous Gamma approximation of site-wise rate variation was left at the default value of 1.

Taking these simulated alignments, we estimated heterogeneous models with *Janus* as described in the main text (e.g., the same search parameters for the empirical data sets: "-ue, -ul, -g"). Across these 400 simulated datasets, we detected zero false positives (no data shown). Thus, our analyses of the sensitivity of *Janus* under a range of tree shapes recovered in the study of the avian molecular phylogeny are consistent with the conservative tendency noted in [5]. Overall, our simulations suggest the signal of molecular model shifts described in the main text is robust to the pattern of rapid cladogenesis observed for Aves and that molecular model shifts are only detected when empirical patterns of sequence variation are strong.

### Assessing the coincidence of model shifts with the K–Pg boundary

Although we detected only a small proportion of assessed nodes to exhibit substitution model shifts (~10%), most of these are detected on nodes for which the K–Pg boundary is included within recently estimated ranges of divergence date uncertainty [1, 6, 7]. In the present case, this is almost exclusively < 5-10 Ma relative to the K–Pg boundary in the MRL3 supertree (Figure 1 and Supplementary Figure 1 a, b). To assess this hypothesis quantitatively, we modeled the Bernoulli probability of a model shift as a function of time distance to the K–Pg boundary, accounting for potential confounding effects of phylogenetic nonindependence, tree shape, and phylogenomic discordance.

We coded a binary dependent variable representing the presence or absence of a novel macroevolutionary regime identified by molecular model shifts and assigned 1/0 to each node using the procedure described below. This strategy is conservative, as multiple shifts in distinct genetic data types often occur along a single edge (e.g., our approach models the minimum number of implied macroevolutionary regimes). We then coded the time distance to the boundary as an independent variable, defined as the stem age for a given focal node minus 66 Ma [by definition, a model shift identified for a given focal node may have occurred anywhere along the branch leading to it]. These values were log-transformed after taking their absolute values, which defines our investigation in terms of proportional proximity to the K—Pg boundary. We fit models of this type with the maximum likelihood 'phyloglm' approach in the *phylolm* R package [10-12], and compared models using the helper functions available at https://github.com/mrhelmus/phylogeny_manipulation. We additionally estimated 1000 bootstrap replicates for each data type to assess the uncertainty around model parameters.

Typically, phylogenetic regressions are applied to data sets associated with contemporary tips on a phylogeny. In our case, we assessed the probability of regime shifts tagged to internal nodes on the phylogeny (e.g., the presence or absence of a model shift in any data type). Because phylogenetic regression techniques use a variance-covariance matrix (VCV) describing shared path lengths proportional to the phylogenetic covariances of trait values [13], these methods are valid for phylogenies with non-contemporaneous tips. We apply this logic and assign molecular model shifts as binary states characterizing negligible-length terminals grafted to each internal node. With this "trick," we can extend phylogenetic regression techniques to evaluate properties or states measured for internal nodes [14]. Our overall regression strategy, therefore, relies on a modified phylogeny that includes 197 non-contemporaneous, negligible-length terminals grafted to each internal node (Supplementary R code).

Lastly, we investigated whether or not patterns of phylogenomic conflict hypothesized to be associated with the K–Pg boundary (e.g., [7, 15]) could confound statistical associations between the probability of a novel molecular model regime and the time distance to the K–Pg boundary. This consideration serves two purposes: 1) we sought to understand how the detection of a molecular model shift may be related to potentially co-varying patterns of phylogenomic discordance, which we also expect to be somewhat correlated with proximity to the K–Pg boundary [15], and 2) we sought to account for the fact that branch lengths estimated from concatenated datasets can contain artifacts derived from model-misspecification related to gene-tree/species tree discordance (e.g. [16]). To address these possibilities, we quantified patterns of phylogenomic discordance across each data type and included a metric of discordance as an additional covariate in logistic regression models. We processed each gene tree (910 separate loci) to collapse nodes with less than 95% ultrafast-bootstrap support [17]. We then coded each node's percentage of discordant gene trees relative to the fixed MRL3 topology for each data type (Supplementary R Code). We applied a variance-stabilizing transformation by taking the arcsine-square root of the percentage discordance values; this transformation recognizes the biological limits on the concept of discordance while spreading out the weight of extreme values. Finally, we excluded extant terminals when running logistic regression models because discordance cannot be measured for these edges. We then compared the results from alternative models considering different levels of phylogenomic discordance as a covariate (Supplementary Figure 1a, b, c, d). We depict alternative analyses considering low (mean – 1 standard deviation), mean, and high (mean + 1 standard deviation) levels of discordance.

#### Results

Phylogenetic logistic regression models show a rapid increase in the probability of a molecular model shift as the time distance to the boundary decreases (Supplementary Figure 1a-d). In all models (except those representing mtDNA), time distance to the boundary is a significant predictor of model shift probability ($p << 0.05$). Considering low (Supplementary Figure 1b), mean (Supplementary Figure 1c), or high (Supplementary Figure 1d) values of phylogenomic discordance has a limited effect on the predicted probability of model shifts and does not appear to strongly confound any associations. Considering the predicted probability of model shifts across the "merged" nuclear data signal at the time of the oldest shifts in the dataset (marked with leader lines; black curves in Supplementary Figure 1), we find a weak inverse relationship between low, mean, and high values of discordance, and model shift probability [63%, 59%, 55%]. This implies a weak but consistent confounding effect, with higher levels of discordance leading to a weaker probability of model shifts. These trends are shared with patterns discordance and model shifts in exon and UTR datasets [42%, 32%, 24%, and 44%, 37%, 30%, respectively]. The signal of shifts in introns, however, shows the opposite trend, with patterns of low, mean, and high discordance leading to a linear increase in shift probability [19%, 26%, and 34%, respectively], implying that the dynamics of model shifts in introns may be different from those in other nuclear data types. In any case, the bootstrapped confidence intervals for different levels of discordance across introns largely overlap, challenging clear interpretations from these data.

In sum, and in general, considering phylogenomic discordance in these models does not reject an inference of an association between time distance to the K–Pg boundary and an increased probability of model shifts. Phylogenomic discordance itself is not a significant covariate in any individual pGLM model, and models which exclude discordance entirely are weakly to moderately preferred under AICc (~ 1.5 AICc unit for the "aggregate" model, < 1 AICc units for exons, ~5 AICc units for introns, and ~1.1 AICc units for UTRs). Notably, while there is some collinearity between time distance to the boundary and discordance, the association is only strong in non-phylogenetic regression models (not shown but see Supplementary R code). Thus, models which account for phylogeny may also largely account for collinearity between discordance and time distance to the boundary.

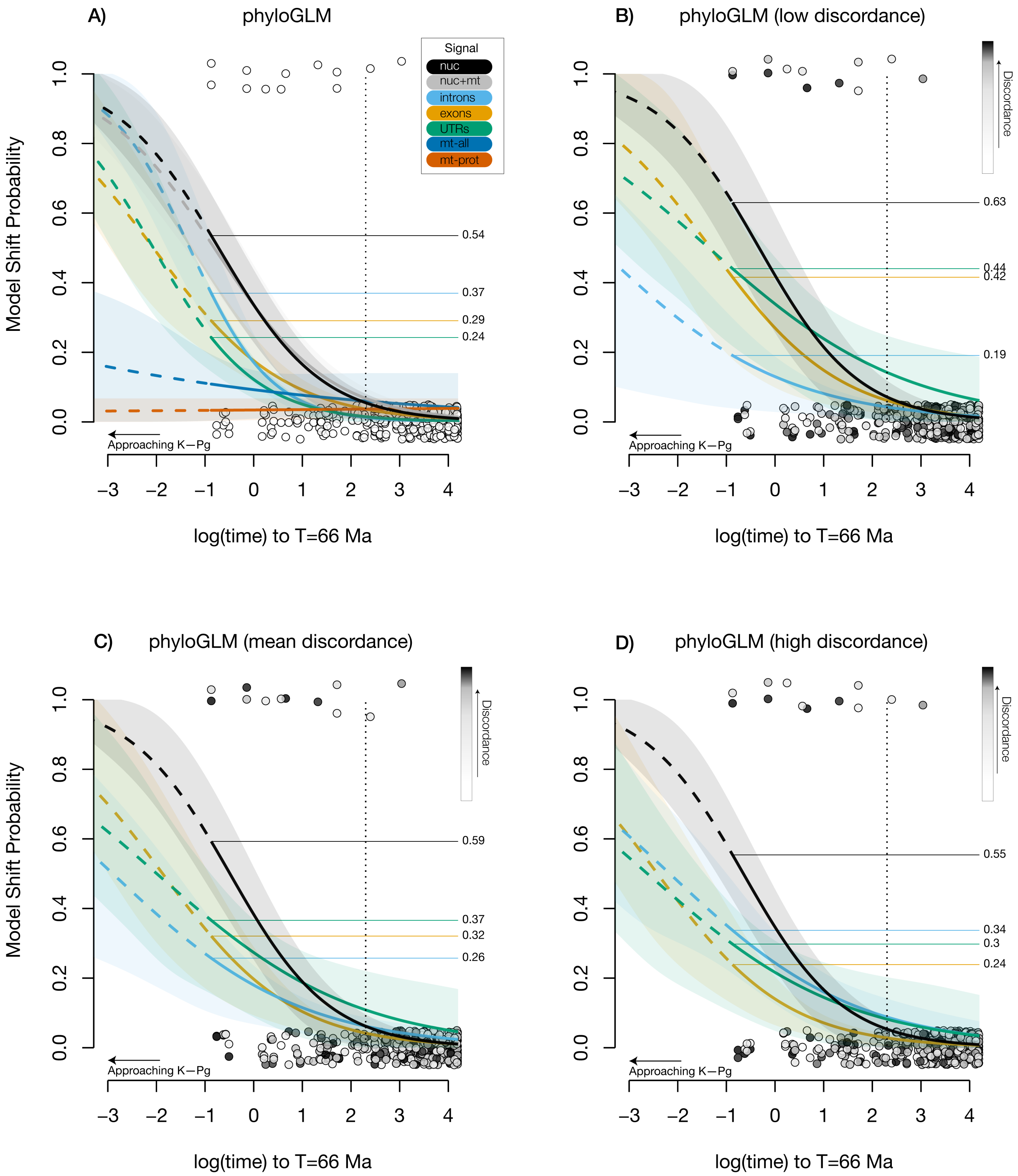
A)
phyloGLM
B)
phyloGLM (low discordance)
C)
phyloGLM (mean discordance)
D)
phyloGLM (high discordance)
Signal
nuc
nuc+mt
introns
exons
UTRs
mt-all
mt-prot
Discordance
Model Shift Probability
0.0
0.2
0.4
0.6
0.8
1.0
0.54
0.37
0.29
0.24
0.63
0.44
0.42
0.19
0.59
0.37
0.32
0.26
0.55
0.34
0.3
0.24
Approaching K−Pg
−3
−2
−1
0
1
2
3
4
log(time) to T=66 Ma

**Supplementary Figure 1. Model shift probability as a function of time distance to the K-Pg boundary.** Binomial probability of novel molecular model regimes as a function of time distance to the K–Pg boundary (e.g., time distance approaches zero, going from right to left, log scale). Logistic regression curves and associated 70% highest density intervals from bootstrap replicates are depicted over the jittered empirical probability distribution for individual nodes (circular markers). Model projections outside the range of the observed data are shown as dashed curves. In panel A, the probability of a model shift in the aggregate signal from nuclear genetic data (black curve) increases to ~54% at the time of the oldest model shift. The probabilities of shifts across each nuclear genetic data type are somewhat lower (~26%-37%). In contrast, the probability of a shift in mtDNA (all data, or proteins only) is not associated with time distance to the boundary (flat curves). This pattern for mtDNA is consistent with the overall low number of shifts inferred across mtDNA genomes ($n < 3$). Panel B-D depict logistic regression curves from models which include phylogenomic discordance as a covariate and show patterns of phylogenomic discordance with a white-to-black color ramp. Discordance is not detected as a significant covariate in any model, and models with discordance have higher AICc values, suggesting they are not a better fit for the data. Nevertheless, comparing model predictions with low, mean, and high discordance values show a very weak confounding effect, with higher levels of discordance leading to a weaker predicted probability of model shifts, except for introns (see Results).

### Functional dimensions of sequence variation

Estimates of the optimal configuration of molecular model shifts suggested equilibrium base frequencies differed substantially across the identified regimes in each data type (Figure 1, Supplementary Figure 7a-d). Although our dataset was not originally conceived to examine functional characteristics of the genome [1], our new assembly and annotation into distinct data types (exons, introns, UTRs, and mtDNAs) presents us with the opportunity to perform a preliminary assessment of functional variation. As noted in [5], compositional shifts may result from non-demographic processes such as selection on codon usage for translation accuracy or even gene expression [18, 19]. Therefore, we estimated several nucleotide-based metrics intended to quantify the degree of codon usage bias. Further, as we generated phased haplotype data, we explored whether patterns of allelic variation in this data set might be related to compositional shifts at the macroevolutionary scale.

For nuclear coding sequences, we evaluated three metrics of codon usage bias: Synonymous Codon Usage Order SCUO [20], Effective Number of Codons $\hat{N}_c$ [21], and a modified version of Effective Number of Codons, or $\hat{N}'_c$ [22]. SCUO measures the nonrandomness in synonymous codon usage and ranges from 0 (totally random) to 1 (totally biased), and is derived from Shannon information theory [23]. $\hat{N}_c$ measures the effective number of codons and ranges from 20 (one codon per AA) to 61 (alternative synonymous codons equally likely). $\hat{N}'_c$ additionally accounts for variation in background nucleotide composition. We, therefore, expected $\hat{N}'_c$ to be the least sensitive to variation in patterns of synonymous codon usage concerning bipartitions identified partly based on compositional variation. Lastly, we assessed nucleotide diversity $\pi$, as approximated by the sum of the branch lengths separating phased alleles. As with life-history traits, we log-transformed metrics of codon bias before comparative phylogenetic analysis. We estimated whether or not variation in these statistics was different across taxon partitions identified by *Janus*, using phylogenetic ANOVA assuming a Brownian model of trait evolution [24, 25], with 10,000 simulations [26] to assess significance.

#### Results

We found no statistical evidence that patterns of nucleotide diversity are different across molecular model shift regimes after accounting for non-independence due to phylogeny (F = 0.933 p = 0.98), though we emphasize that this is an exploratory analysis in the absence of population-level sampling at the species level. Next, we found that patterns of synonymous codon usage may be significantly different across model regimes in the omnibus ANOVA, and for many cases of post-hoc comparison after correction for multiple hypothesis tests (SCUO; F: 17.6, p = 0.005, $\hat{N}_c$; F: 17.23, p = 0.005, $\hat{N}'_c$ F: 27.23, p = 0.0003). These results are depicted in Supplementary Figure 2. Notably, the $\hat{N}'_c$ metric appears to be the most sensitive to patterns of variation in codon usage, identifying many cases in which post-hoc pairwise comparisons are statistically different. For example, with $\hat{N}'_c$ the unnamed sister clade to Tinamiformes (a clade uniting Rheiformes, Casuariiformes, and Apterygiformes) appears to have the strongest signal of differentiation, with 5/7 comparisons implying significantly different patterns of codon usage (Supplementary Figure 2), sans the groups Passeri or Tinamiformes). In sum, for all eight groups identified by *Janus* as occupying distinct molecular model regimes in exons, we find evidence of significant differences in patterns of codon usage in comparison to at least one other candidate group. Although it was not our focus here, these results strongly imply that identifying compositional shifts with methods like *Janus* may be fruitful in generating hypotheses about functional variation in coding sequences.

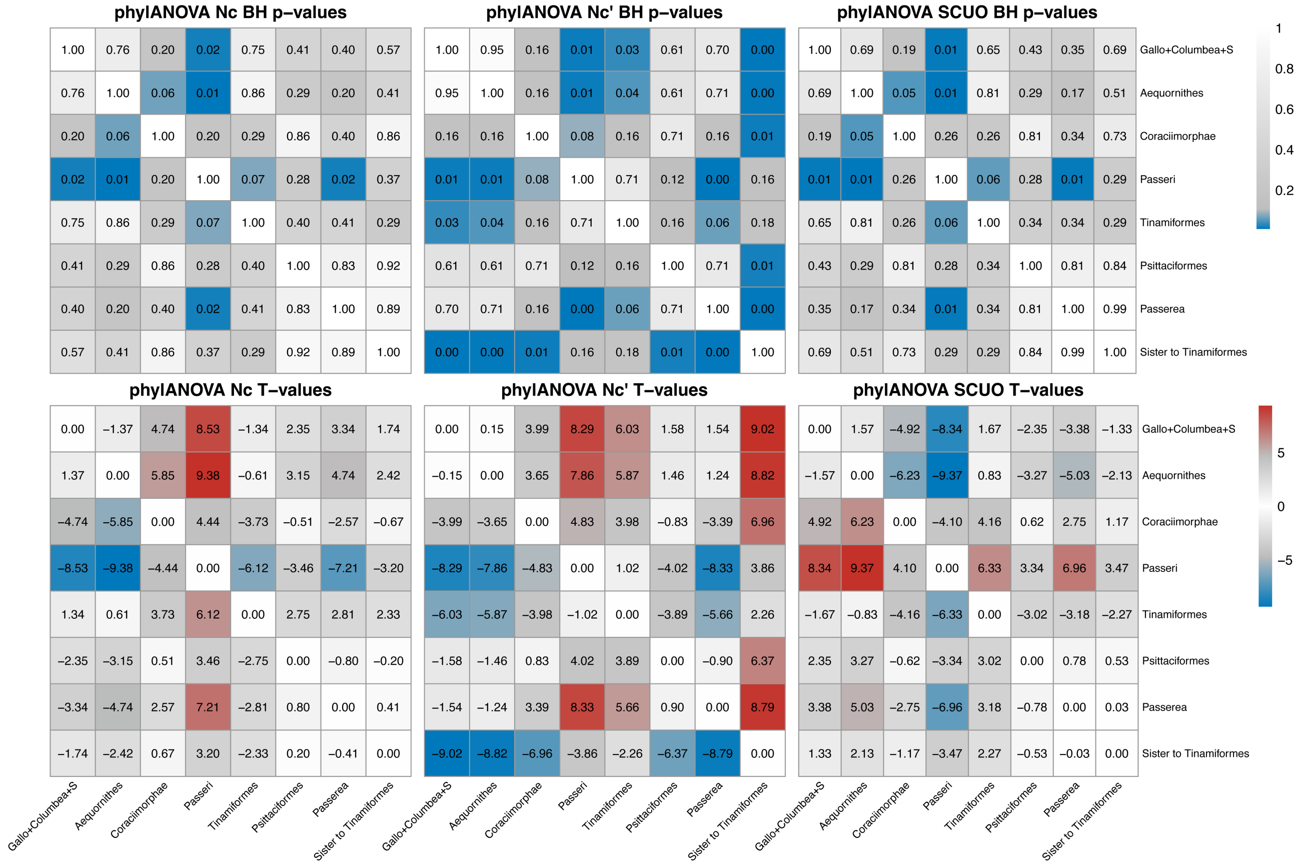


**Supplementary Figure 2.** Synonymous codon usage patterns are associated with molecular model shifts. For exon data, we evaluated whether patterns of synonymous codon usage differed across taxon partitions identified in the analysis of molecular model shifts. In the top row, we depict matrices of BH-corrected p-values for post-hoc pairwise comparisons from phylogenetic ANOVA. In the bottom row, we depict T-values for each respective test. See the Supplementary Appendix for additional detail.

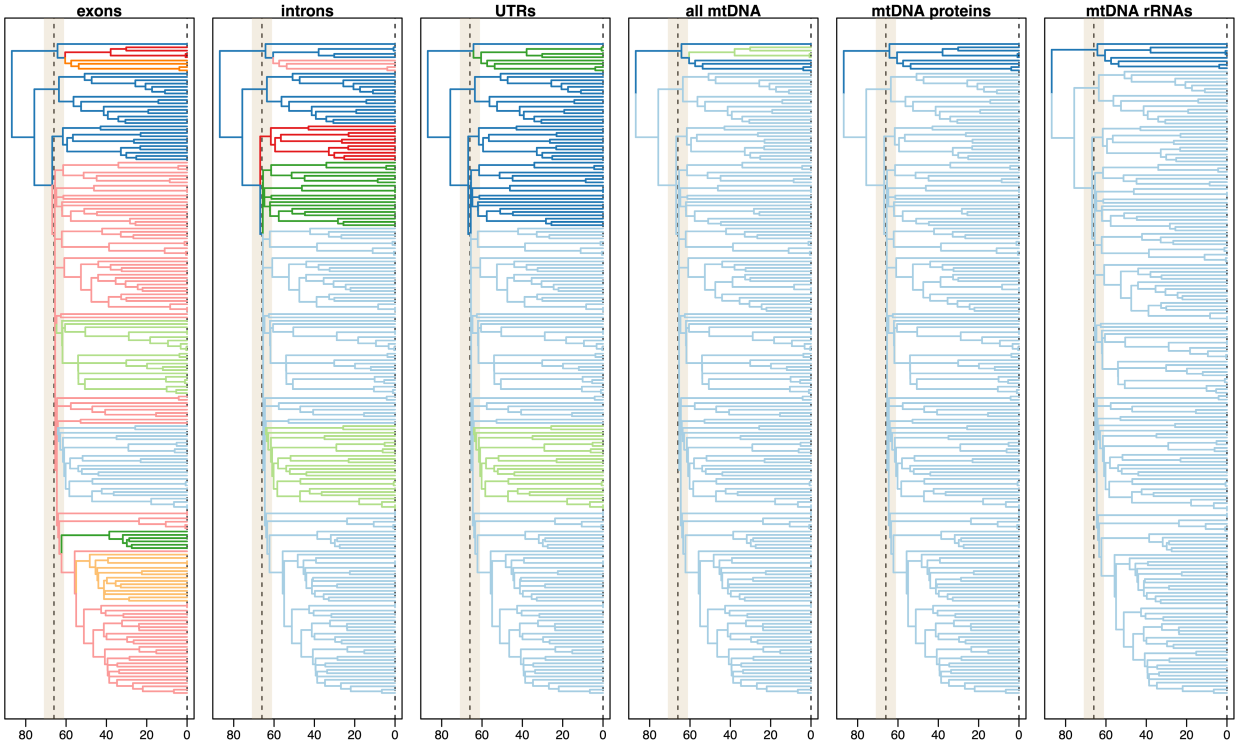


**Supplementary Figure 3.** Patterns of molecular model shifts for each evaluated data type. Changes in branch color indicate model shifts. Figure 1 contains this information in aggregate. Model shifts are most common in exon data, followed by introns, UTRS, and mtDNAs. We note that two model shifts are detected in the analysis of the whole mtDNA dataset (Neognathae and Tinamiformes), whereas analysis of mtDNA proteins or rRNAS separately identifies the same model shift (Neognathae only). We interpret this difference to reflect increased power from the combined mtDNA dataset. Also, see Supplementary Figure 7a-d for an expanded view.

### Assessing the coincidence of molecular shifts and shifts in θ(t)

As described in the main text, we assessed statistical support for the hypothesis that shifts in θ(t) may have co-occurred with molecular model shifts using a bootstrapping approach implemented in ℓ1ou [27]. Under models relying on the AICc criterion, median support was 87.2% [0.59-0.91] for shifts across the eight-dimensional dataset (pie charts in Figure 1, Supplementary Figure 2). Considering the more conservative pBIC criterion, median support was modestly lower: 76.2% [0-97.3]. For the latter case, three candidate edges received no support relative to the false positive rate: the edge leading to Oscine Passeriformes and the two edges reflecting the stem of Otidae or its unnamed sister clade. Notably, these latter two edges reflect the two deepest splits in Passerea [7], while the edge leading to the MRCA of Passerea receives the strongest relative support under pBIC (97.3%). Applying Fisher's exact test to these inferences supports all cases except those noted above, for which a test of equality in proportion is not rejected at $p = 0.05$. All other cases are supported at $p < 10^{-4}$ under both AICc and pBIC except for two cases: Notopalaeognathae (AICc: $p = 0.238$, OR = 1.49; pBIC: $p$

= 0.04, OR = 2.1), and the sister clade to Tinamiformes (AICc: $p$ = 0.015, OR = 2.19, pBIC: $p$ < $10^{-4}$, OR = 4.7) (Supplementary Figure 4, Supplementary Table 2).

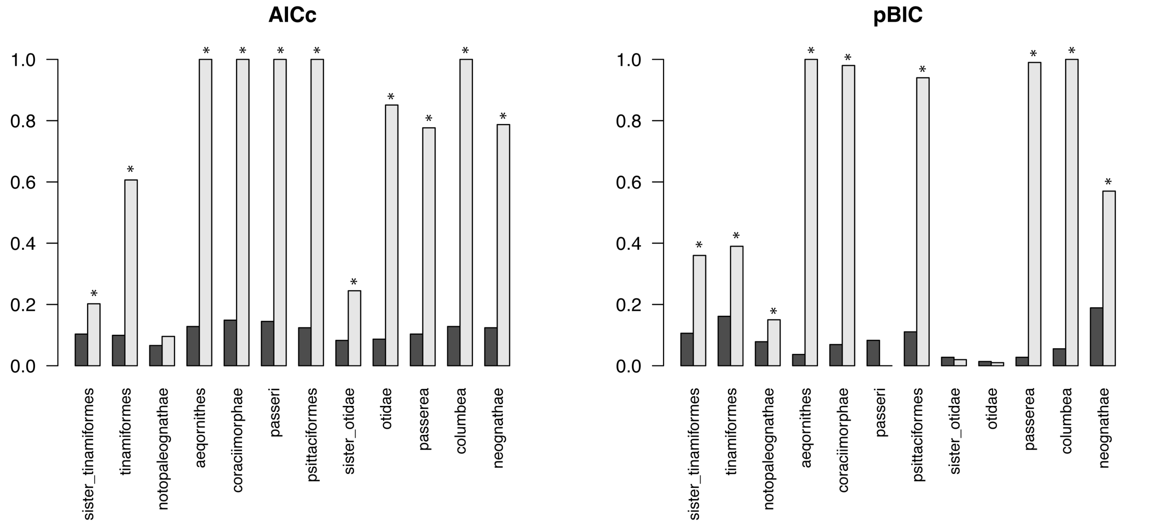


**Supplementary Figure 4.** Relative support for shifts in trait optima θ(t). For each criterion (AICc or pBIC), we depict the frequency of empirical positive shift detections from ℓ1ou bootstrapping (light gray) relative to the frequency of false positive detections (dark gray) generated with a dataset simulated under multivariate Brownian motion (e.g., no shifts in θ). Cases in which the empirical positive detection rate is significantly greater than the false positive rate (Fisher's exact test) at p=0.05 are marked with an asterisk (see supplementary methods and Supplementary Table 2).

### Assessing LHT importance with a Random Forest Classifier

To identify which, if any, LHTs (life-history traits) may be good predictors of molecular model shifts, we used an approach from the field of supervised machine learning known as Random Forests [28, 29]. The Random Forests approach generates a classification model based on a population of decision trees [30], and can naturally assess the relative importance of different features with high accuracy [31]. Here, we focus on the classification of taxon partitions (groups of terminals) identified in analyses of exon data, though alternative analyses of taxon partitions identified in other data types generated similar results (not shown). Although it may be possible to incorporate aspects of phylogenetic distance into these analyses [31, 32], non-parametric machine learning methods like Random Forest make no assumptions about the distribution of the underlying data and can handle skewed or multi-modal data as well as categorical data; thus, accounting for phylogenetic non-independence in the data is not required in the same way as when conducting, for example, a GLS analysis.

First, we split our life-history data into training and test datasets with a 70/30 split, accounting for stratified sampling. We then used *tidymodels* [33] to build a recipe for data preprocessing, specifying several steps: 1) removing any variables correlated with others at a Pearson correlation coefficient > 0.95, 2) normalizing (centering and scaling), 3) creating dummy variables for categorical variables with one hot encoding, and 4) generating synthetic positive instances using ADASYN algorithm [34] to increase the sample size of small groups to at least 50% of the size of the largest group (setting the number of neighbors to 2).

Next, we specified the structure of the model, including hyperparameters "mtry" (number of features to sample, set to tune automatically) and "min_n" (minimum number of data points in a node to allow further splitting, set to tune automatically). We set the number of trees in a forest to 1000 and specified the "randomForest" engine [35]. To tune the hyperparameters, we used k-fold cross-validation with ten folds repeated ten times. We selected the best model from the hyperparameter tuning based on the area under the Receiver Operating Characteristic Curve (AUC) using the estimator from [36], and fit it to the training set. The AUC can be interpreted as the probability that a randomly chosen positive example is ranked above a randomly chosen negative example and ranges from 0 to 1, with values closer to 1 reflecting better model performance. Lastly, we estimated permutation-based variable importance [37] using the VIP R package [38] with 500 simulations. See supplementary R script RandomForest_var_imp.R for details.

#### Additional models of life-history traits

In the main text, we present visualizations of OUM models in which trait optima $\theta(t)$ can change across specified regimes (Figure 2, right); in this case, macroevolutionary regimes are defined *a priori* on the basis of molecular model shifts. These results stem from analysis of nuclear genetic data, emphasizing molecular changes associated with the radiation of Neoaves. Indeed, for most cases within this superordinal clade, we observe decreases in the estimates for $\theta(t)$ associated with molecular model shifts relative to the ancestral regime $\theta_{anc}$. Based on our Random Forests model, we visualized two features with the highest relative variable importance scores ("ChickPC1'' and "adult body mass," Figure 2, Right). Notably, this set of taxon partitions from *Janus* includes a plesiomorphic group, uniting our single lineage of *Struthio* with the clade Galloanserae on the same molecular regime (e.g., Supplemental Figure 7a). To explore these patterns further, we fit additional OUM models, placing *Struthio* on a separate regime along with the tree's root. This configuration reflects the aggregate signal of taxon partitions identified across nuclear and mitochondrial datasets (Figure 1, e.g., adding the shift detected in mitochondrial data for Neognathae). By placing the large-bodied *Struthio* lineage on a separate regime, we can estimate how its closest sister clades may vary in $\theta(t)$, relative to the ancestral regime, without being potentially confounded by the inclusion of Galloanseres. For these analyses, we re-estimated OUM models for all numerical life-history traits (Supplementary Figure 6, also see Supplementary Figure 5).

Lastly, [39] describes how the analysis of principal components can, under some circumstances, lead to misleading phylogenetic inferences. Specifically, [39] describes a phenomenon by which ordination partitions the variance of a multivariate trait such that the major component axes are biased toward "early burst" patterns. As one of the features we examine is a principal component of multivariate developmental mode data described by [40], we checked for this bias by fitting univariate EB, OU, and BM models to each of the eight examined life-history traits and then comparing model fit with AIC weights. We performed these checks using the *fitContinuous* function in [24] and further validated the results with *OUwie* [41].

#### Results

Placing *Struthio*+root on a separate macroevolutionary regime reveals more complex dynamics within the Palaeognathae (Supplementary Figure 6). The *Struthio*+root regime is detected to have a much larger $\theta(t)$, magnifying the apparent shift in life-history trait $\theta(t)$ for all other taxon partitions identified by molecular model shift analysis. In this configuration, all candidate regimes identified by *Janus* show reduced $\theta(t)$ for ChickPC1 or adult body mass relative to the ancestral regime. We observe similar patterns for clutch size, and a somewhat weaker signal for generation length (Supplementary Figure 5, 6). These analyses corroborate patterns revealed by Random Forest variable importance analysis, which proposed ChickPC1 and adult body mass as most strongly associated with molecular model shifts. Given the challenges in accurately estimating evolutionary parameters like $\theta(t)$ from small sample sizes, we present these additional analyses as exploratory. Nonetheless, all estimates are consistent with patterns described in the main text. Further, checks for a bias toward early burst patterns in the data did not reveal any life-history traits for which "EB" was selected as the optimal model. "OU" models carried nearly 100% of the model weight in all cases (OU > BM > EB in all cases; see Supplementary R code). Thus, at the scale of the examined phylogeny, we are not concerned that ordination (in the case of developmental mode data from [40]) has biased these data toward an early burst pattern.

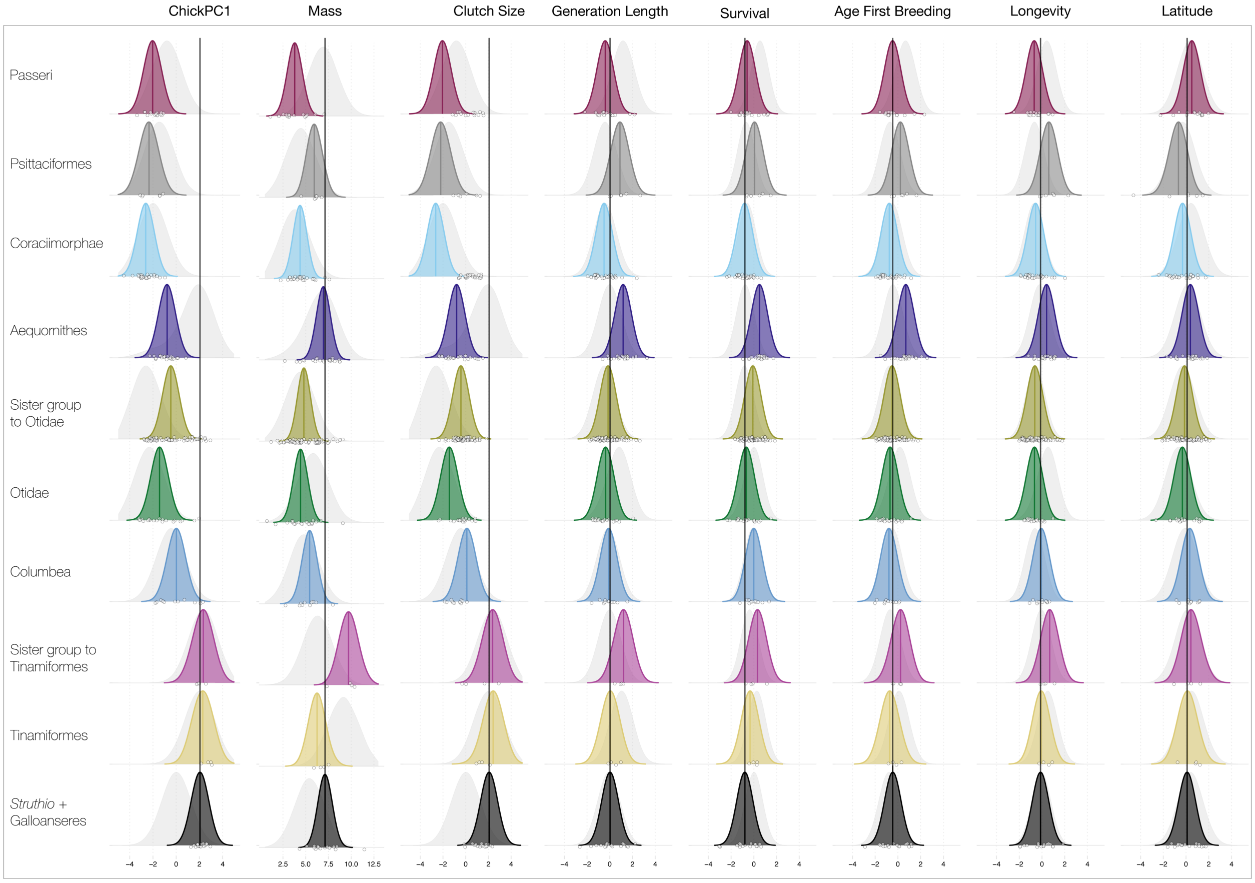


**Supplementary Figure 5.** Complete set of θ(t) shift models (nuclear data configuration). We estimated shifts in θ(t) under an OUM model in OUwie [41]. We describe the results for ChickPC1 and adult body mass in the main text, as these two features are identified as being most important with a Random Forest classifier (See main text and supplementary appendix). Here, we show the complete set of models for each of the eight continuous life-history traits. Colored distributions reflect parametric bootstrap estimates of θ(t), while grayed-out background distributions reflect a summary of 10,000 simulations of tip values from the model (e.g., time = 0). Notably, analysis of clutch size reveals similar patterns to those observed for ChickPC1 and adult body mass (namely, a reduction in the estimated values for θ(t) for the ancestral regime for many Neoavian clades. Although some fluctuations are observed for other traits, none are as pronounced as for ChickPC1 and adult body mass.

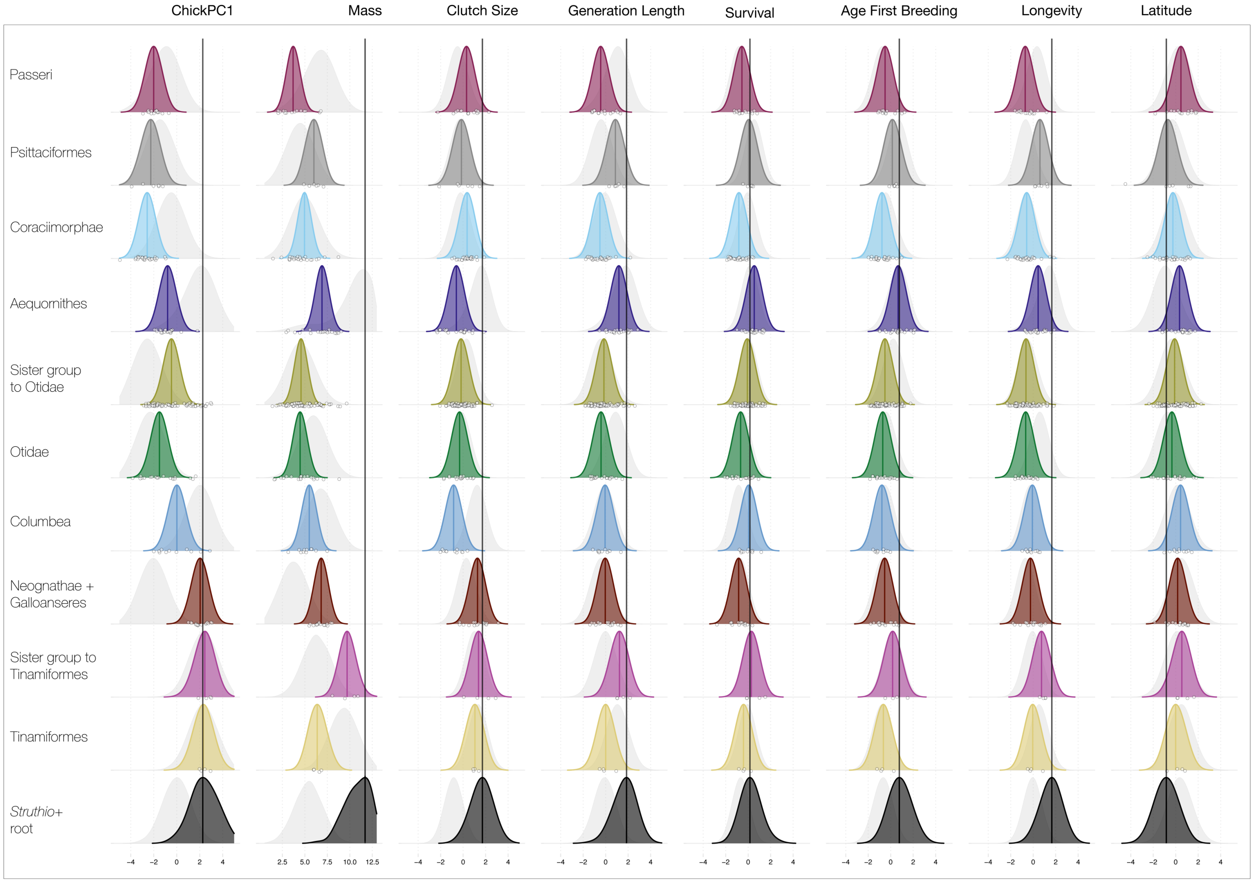


**Supplementary Figure 6.** Complete set of θ(t) shift models (nuc+mt data configuration). We estimated shifts in θ(t) under an OUM model in OUwie [41]. We present ChickPC1 and adult body mass results in the main text, as these two features are identified as most important in a Random Forest classifier (See main text and supplementary appendix). Colored distributions reflect parametric bootstrap estimates of θ(t), while grayed-out background distributions reflect a summary of 10,000 simulations of tip values from the model (e.g., time = 0). Here, we show the complete set of models for each of eight continuous life-history traits– with an essential difference from the results presented in Figure 2 and Supplementary 5. Here, an additional taxon partition is included, reflecting a unique molecular model shift identified in the analysis of mtDNA (Neognathae). This has the effect of placing *Struthio* on a regime separated from Galloanseres (e.g., Figure 2, Supplementary Figure 5). The Galloanseres are thus united with a regime including the Neognathae MRCA. Shift patterns for ChickPC1 and clutch size θ(t) are essentially identical to those presented in prior analyses – however, investigation of body mass reveals more complex dynamics within the Palaeognathae. The *Struthio* regime contains the tree's root and is naturally estimated to have a much larger θ(t). This, in turn, induces an apparent decrease in θ(t) for body mass in Tinamiformes and more substantial decreases in θ(t) for all other cases. As noted in the supplementary results, these patterns suggest that *all molecular model shifts* (either nuclear or mtDNA) may indicate derived decreases in body mass or ChickPC1 θ(t). Similar fluctuations are observed for other traits (e.g., generation length), but none are as pronounced as the patterns for ChickPC1 and adult body mass.

Supplementary Figure 7a

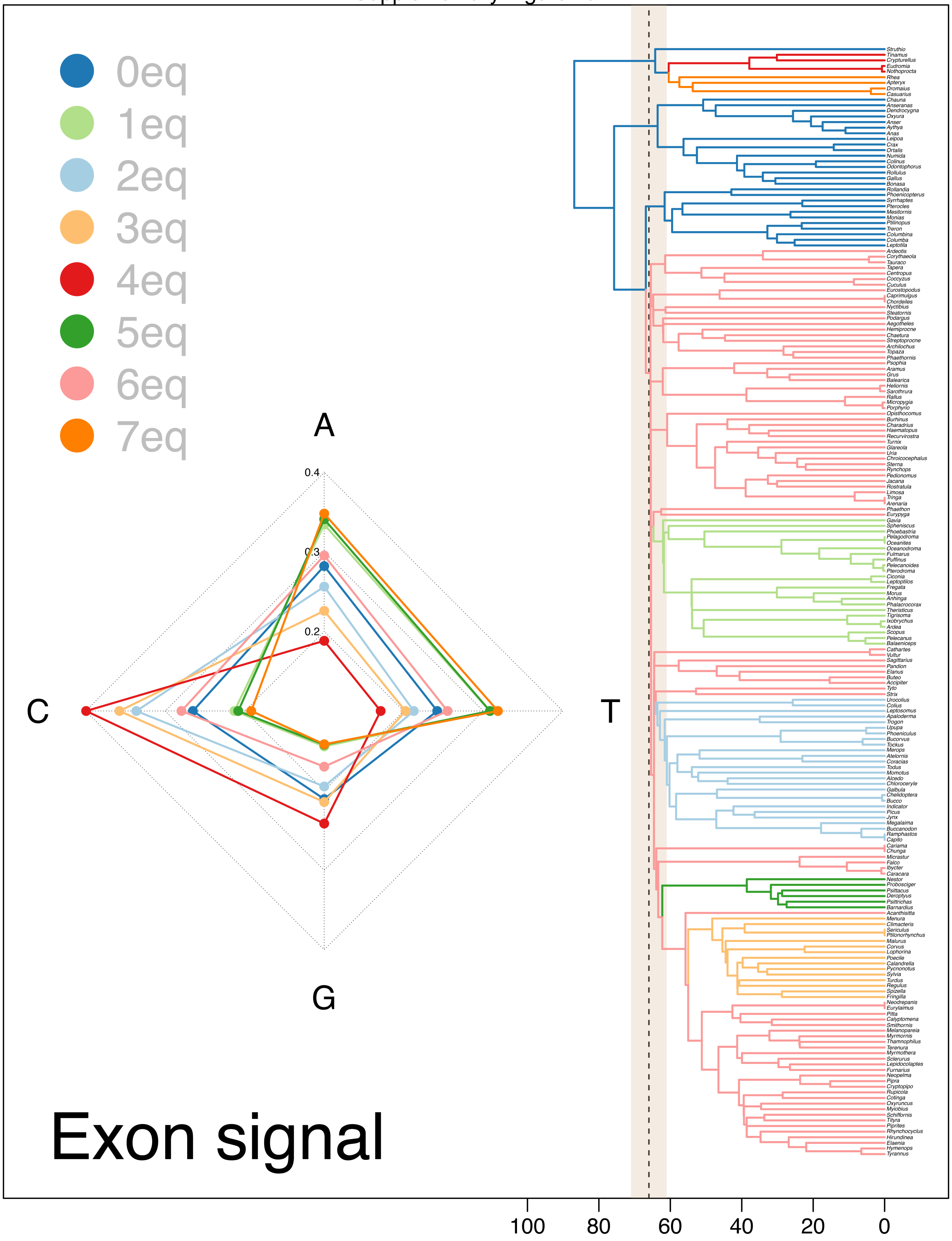

Supplementary Figure 7b

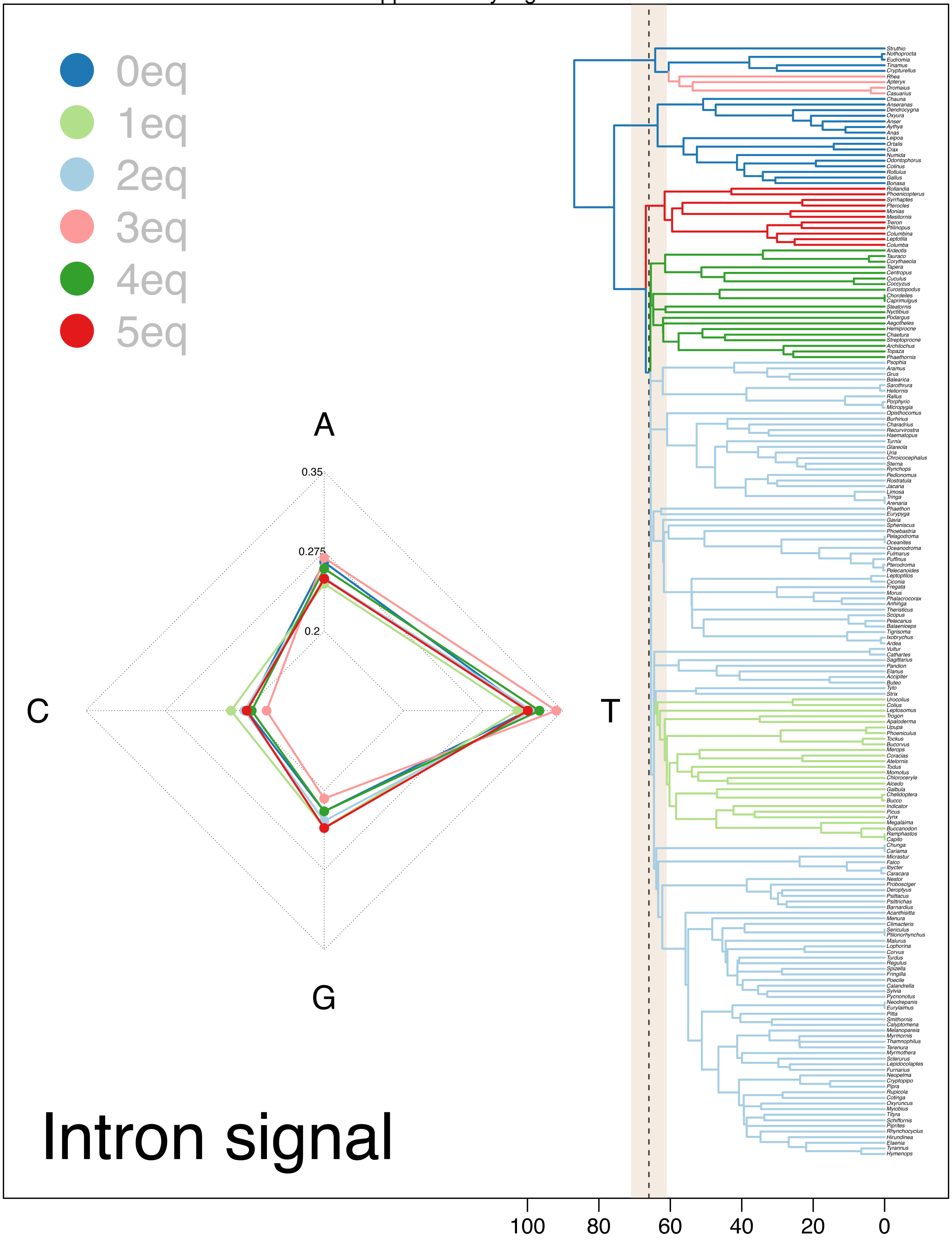

Supplementary Figure 7c

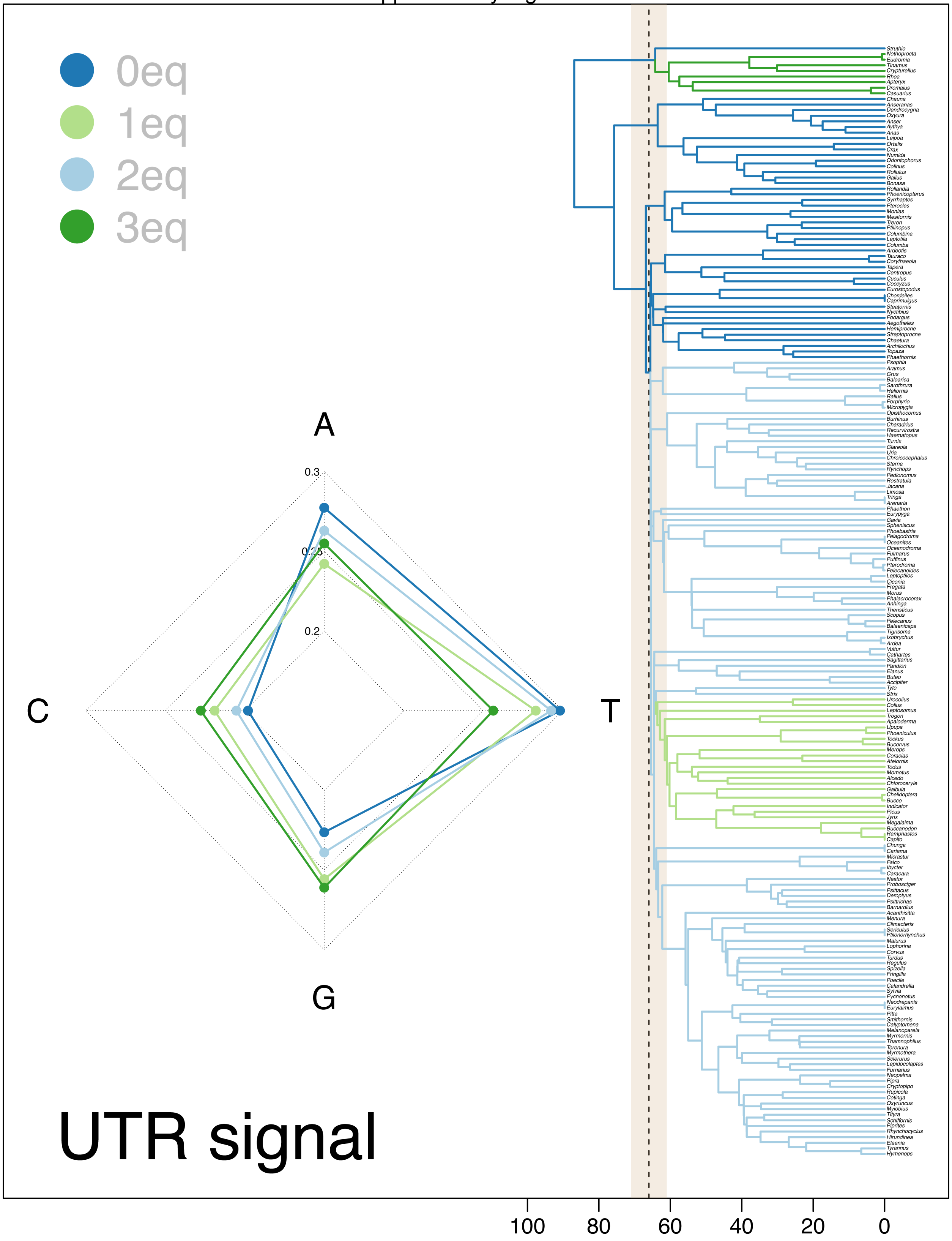

Supplementary Figure 7d

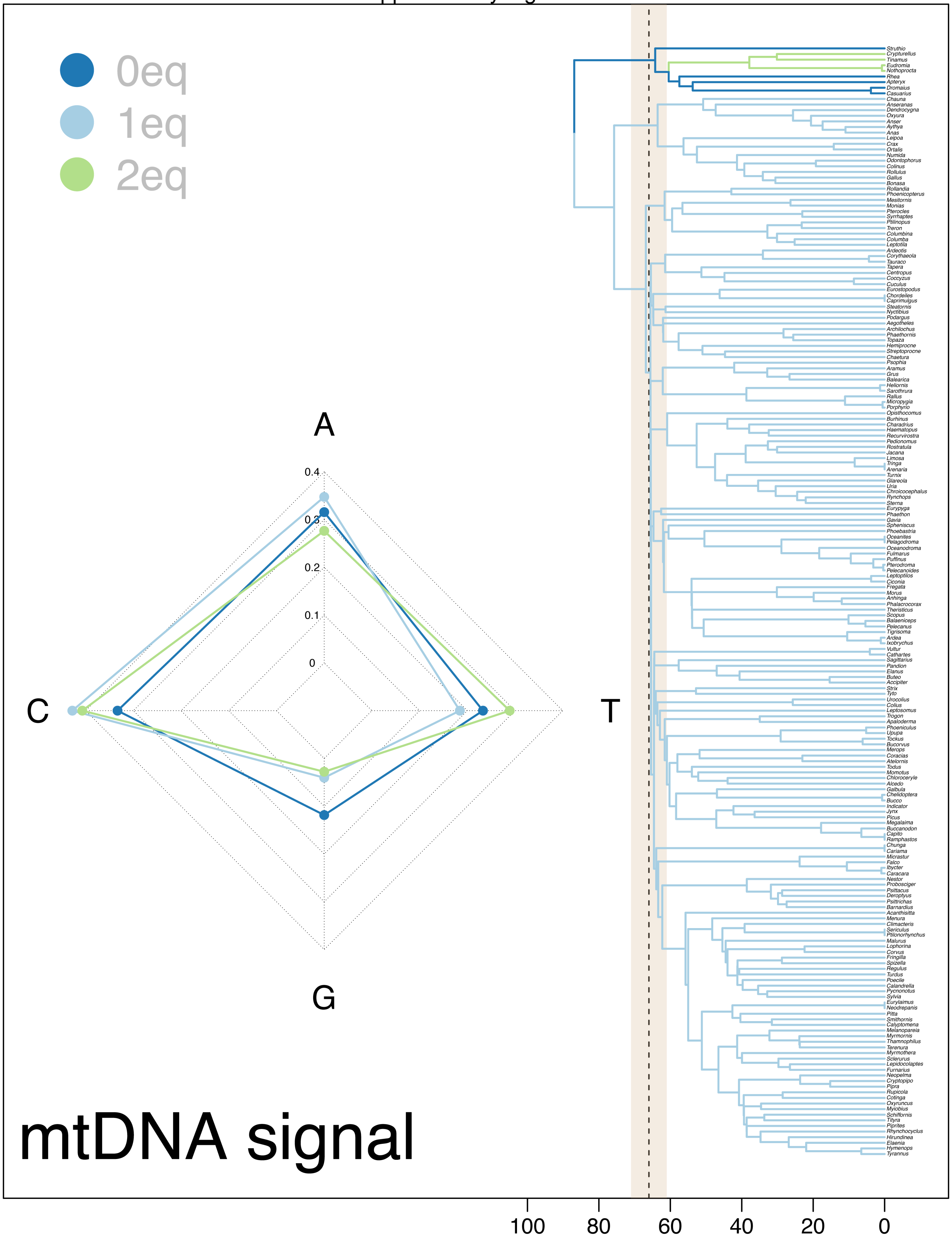

**Supplementary Figure 7a-d.** Maps of estimated equilibrium base frequencies for each taxon partition identified by *Janus* in analyses of different genetic data types. For each plot a-d, we show a radar plot depicting the relative estimated equilibrium base frequencies for each taxon partition. Colors match those indicated in the legend and correspond to distinct phylogenetic regimes. As noted in the main text and references to Figure 1, the most significant deviations from empirical base frequencies are identified in our exon dataset (see main text discussion) and are concentrated around a brief interval shortly following the end-Cretaceous mass extinction. All molecular model shifts, except for a shift in exon data in Oscine Passeriformes and mtDNA for Neognathae are plausibly associated with lineage diversification near the K–Pg boundary (e.g., Supplementary Figure 1).

| Clade | Authority | Exons | Introns | UTRs | mtDNAs | $\beta_0$ | $\beta_0$ 95% HPD | $\beta_{mass}$ | $\beta_{mass}$ 95% HPD | *pp* | Power law |
|---|---|---|---|---|---|---|---|---|---|---|---|
| Aves | Linneaus, 1758 | anc. | anc. | anc. | anc. | -3.40 | -5.68, -1.79 | 0.69 | 0.52, 0.86 | - | $0.03M^{0.69}$ |
| Paleognathae | Pycraft, 1900 | - | - | - | - | - | - | - | - | - | - |
| Notopaleognathae | Yuri et al. 2013 | - | - | x | - | -3.82 | -6.86, -0.01 | 0.73 | 0.5, 0.89 | 0.06 | $0.02M^{0.73}$ |
| Tinamiformes | Huxley, 1872 | x | - | - | x | -3.92 | -4.99, -2.8 | 0.76 | 0.6, 0.94 | 0.10 | $0.02M^{0.76}$ |
| Rheiformes, Casuariiformes, and Apterygiformes | Unnamed clade | x | x | - | - | -4.32 | -5.65, -2.83 | 0.77 | 0.62, 0.91 | 0.15 | $0.01M^{0.77}$ |
| Neognathae | Pycraft, 1900 | - | - | - | x | -3.66 | -4.71, -2.93 | 0.72 | 0.61, 0.87 | 0.02 | $0.03M^{0.72}$ |
| Columbea | Jarvis 2014 | - | x | - | - | -4.28 | -4.97, -3.49 | 0.84 | 0.71, 0.98 | 0.39 | $0.01M^{0.84}$ |
| Passerea | Jarvis 2014 | x | - | - | - | -3.32 | -6.82, -0.05 | 0.69 | 0.51, 0.9 | 0.07 | $0.04M^{0.69}$ |
| Otidae | Wagler 1830 | - | x | - | - | -3.47 | -3.77, -3.15 | 0.68 | 0.61, 0.74 | 0.12 | $0.03M^{0.68}$ |
| Reminader of Neoaves | Unnamed clade | - | x | x | - | -3.27 | -3.32, -2.93 | 0.68 | 0.62, 0.7 | 0.38 | $0.04M^{0.68}$ |
| Aequornithes | Jarvis 2014 | x | - | - | - | -3.53 | -3.98, -2.95 | 0.73 | 0.65, 0.79 | 0.08 | $0.03M^{0.73}$ |
| Coraciimorphae | Sibley & Ahlquist, 1990 | x | x | x | - | -3.86 | -4.13, -3.61 | 0.75 | 0.7, 0.8 | 0.98 | $0.02M^{0.75}$ |
| Psittaciformes | Wagler, 1830 | x | - | - | - | -3.84 | -4.71, -2.89 | 0.75 | 0.6, 0.91 | 0.11 | $0.02M^{0.75}$ |
| Passeri | Linnaeus, 1758 | x | - | - | - | -3.17 | -3.64, -2.71 | 0.65 | 0.52, 0.79 | 0.04 | $0.04M^{0.65}$ |

**Supplementary Table 1.** Summary of molecular shifts across avian higher taxa. For each clade on which we detect a molecular model shift, we note the taxonomic authority and which data types are identified to have model shifts (with an 'x'). We report the associated metabolic allometric parameter estimates from *bayou* for the reader's reference: intercept ($\beta_0$), slope ($\beta_{mass}$), their associated 95% highest posterior density intervals (HPD), the posterior probability of an allometric shift occurring on a particular edge (*pp*), and the estimated power law describing the association between metabolic rate and body mass ($e^{\beta_0} \times mass^{\beta_{mass}}$). Notably, the diverse clade Coraciimoraphae is detected to have molecular model shifts in all nuclear genetic data types and maximal posterior probability for a shift in metabolic allometry.

| **AICc** | Authority | p-value | 2.50% | 97.50% | Odds Ratio |
|---|---|---|---|---|---|
| Aves | Linneaus, 1758 | anc. | anc. | anc. | anc. |
| Paleognathae | Pycraft, 1900 | - | - | - | - |
| Notopaleognathae | Yuri et al. 2013 | 0.24 | 0.65 | Inf | 1.49 |
| Tinamiformes | Huxley, 1872 | 0.00 | 8.15 | Inf | 13.84 |
| Rheiformes, Casuariiformes, and Apterygiformes | Unnamed clade | 0.01 | 1.20 | Inf | 2.19 |
| Neognathae | Pycraft, 1900 | 0.00 | 14.74 | Inf | 25.74 |
| Columbea | Jarvis 2014 | 0.00 | 201.35 | Inf | Inf |
| Passerea | Jarvis 2014 | 0.00 | 16.78 | Inf | 29.64 |
| Otidae | Wagler 1830 | 0.00 | 30.77 | Inf | 58.60 |
| Reminader of Neoaves | Unnamed clade | 0.00 | 1.96 | Inf | 3.58 |
| Aequornithes | Jarvis 2014 | 0.00 | 201.35 | Inf | Inf |
| Coraciimorphae | Sibley & Ahlquist, 1990 | 0.00 | 164.74 | Inf | Inf |
| Psittaciformes | Wagler, 1830 | 0.00 | 201.35 | Inf | Inf |
| Passeri | Linnaeus, 1758 | 0.00 | 169.99 | Inf | Inf |

| **pBIC** | Authority | p-value | 2.50% | 97.50% | Odds Ratio |
|---|---|---|---|---|---|
| Aves | Linneaus, 1758 | anc. | anc. | anc. | anc. |
| Paleognathae | Pycraft, 1900 | - | - | - | - |
| Notopaleognathae | Yuri et al. 2013 | 0.04 | 1.04 | Inf | 2.07 |
| Tinamiformes | Huxley, 1872 | 0.00 | 2.03 | Inf | 3.31 |
| Rheiformes, Casuariiformes, and Apterygiformes | Unnamed clade | 0.00 | 2.76 | Inf | 4.72 |
| Neognathae | Pycraft, 1900 | 0.00 | 3.41 | Inf | 5.43 |
| Columbea | Jarvis 2014 | 0.00 | 469.25 | Inf | Inf |
| Passerea | Jarvis 2014 | 0.00 | 504.78 | Inf | 2945.38 |
| Otidae | Wagler 1830 | 0.78 | 0.03 | Inf | 0.72 |
| Reminader of Neoaves | Unnamed clade | 0.78 | 0.10 | Inf | 0.72 |
| Aequornithes | Jarvis 2014 | 0.00 | 679.55 | Inf | Inf |
| Coraciimorphae | Sibley & Ahlquist, 1990 | 0.00 | 172.19 | Inf | 634.71 |
| Psittaciformes | Wagler, 1830 | 0.00 | 53.72 | Inf | 122.13 |
| Passeri | Linnaeus, 1758 | 1.00 | 0.00 | Inf | 0.00 |

**Supplementary Table 2.** Summary of Fisher's exact test for shifts in life-history trait optima associated with molecular model shifts. With $\ell$1ou, we used parametric bootstrapping to estimate the frequency of shift detections and the null false positive rate for each candidate edge under simulated multivariate Brownian motion. We use Fisher's test to ask if the proportion of empirical shift detections in the bootstrapped dataset is higher than that of null false positives in the simulated dataset. All candidate edges are supported under AICc or pBIC criteria, but not both in a few cases.

**Supplementary Table 3**
Details of mtDNA dataset assembly. See supplemental data at https://github.com/jakeberv/avian_molecular_shifts.

**Supplementary Table 4**
Life-history and metabolic rate datasets. See supplemental data at https://github.com/jakeberv/avian_molecular_shifts/tree/main/LHT_DATA.

### Literature Cited


1. Prum, R.O., et al., *A comprehensive phylogeny of birds (Aves) using targeted next-generation DNA sequencing.* Nature, 2015. **526**(7574): p. 569-573.

2. Alfaro, M.E., et al., *Nine exceptional radiations plus high turnover explain species diversity in jawed vertebrates.* Proceedings of the National Academy of Sciences, 2009. **106**(32): p. 13410-13414.

3. Mitov, V., K. Bartoszek, and T. Stadler, *Automatic generation of evolutionary hypotheses using mixed Gaussian phylogenetic models.* Proceedings of the National Academy of Sciences, 2019. **116**(34): p. 16921-16926.

4. Schwarz, G., *Estimating the dimension of a model.* The annals of statistics, 1978. **6**(2): p. 461-464.

5. Smith, S.A., N. Walker-Hale, and C. Parins-Fukuchi, *Compositional shifts associated with major evolutionary transitions in plants.* bioRxiv, 2022: p. 2022.06.13.495913.

6. Claramunt, S. and J. Cracraft, *A new time tree reveals Earth history's imprint on the evolution of modern birds.* Science Advances, 2015. **1**(11).

7. Jarvis, E.D., et al., *Whole-genome analyses resolve early branches in the tree of life of modern birds.* Science, 2014. **346**(6215): p. 1320-1331.

8. Minh, B.Q., et al., *IQ-TREE 2: New Models and Efficient Methods for Phylogenetic Inference in the Genomic Era.* Molecular Biology and Evolution, 2020. **37**(5): p. 1530-1534.

9. Ly-Trong, N., et al., *AliSim: A Fast and Versatile Phylogenetic Sequence Simulator for the Genomic Era.* Molecular Biology and Evolution, 2022. **39**(5).

10. Tung Ho, L.s. and C. Ané, *A Linear-Time Algorithm for Gaussian and Non-Gaussian Trait Evolution Models.* Systematic Biology, 2014. **63**(3): p. 397-408.

11. Ives, A.R. and T. Garland, Jr., *Phylogenetic Logistic Regression for Binary Dependent Variables.* Systematic Biology, 2010. **59**(1): p. 9-26.

12. Paradis, E. and J. Claude, *Analysis of Comparative Data Using Generalized Estimating Equations.* Journal of Theoretical Biology, 2002. **218**(2): p. 175-185.

13. Harmon, L., *Phylogenetic comparative methods: learning from trees. Self published under a CC-BY-4.0 license.* 2018.

14. Ané, C., *phyloglm troubleshooting*, J. Berv, Editor. 2021, github: github.com.

15. Parins-Fukuchi, C., G.W. Stull, and S.A. Smith, *Phylogenomic conflict coincides with rapid morphological innovation.* Proceedings of the National Academy of Sciences, 2021. **118**(19): p. e2023058118.

16. Mendes, F.K. and M.W. Hahn, *Gene Tree Discordance Causes Apparent Substitution Rate Variation.* Systematic Biology, 2016.

17. Minh, B.Q., M.A.T. Nguyen, and A. von Haeseler, *Ultrafast Approximation for Phylogenetic Bootstrap.* Molecular Biology and Evolution, 2013. **30**(5): p. 1188-1195.

18. Clément, Y., et al., *Evolutionary forces affecting synonymous variations in plant genomes.* PLOS Genetics, 2017. **13**(5): p. e1006799.

19. Qiu, S., et al., *Patterns of Codon Usage Bias in Silene latifolia.* Molecular Biology and Evolution, 2010. **28**(1): p. 771-780.

20. Wan, X.-F., et al., *Quantitative relationship between synonymous codon usage bias and GC composition across unicellular genomes.* BMC Evolutionary Biology, 2004. **4**(1): p. 19.

21. Wright, F., *The 'effective number of codons' used in a gene.* Gene, 1990. **87**(1): p. 23-29.

22. Novembre, J.A., *Accounting for Background Nucleotide Composition When Measuring Codon Usage Bias.* Molecular Biology and Evolution, 2002. **19**(8): p. 1390-1394.

23. Shannon, C.E., *A mathematical theory of communication.* The Bell system technical journal, 1948. **27**(3): p. 379-423.

24. Harmon, L.J., et al., *GEIGER: investigating evolutionary radiations.* Bioinformatics, 2008. **24**(1): p. 129-131.

25. Revell, L.J., *phytools: an R package for phylogenetic comparative biology (and other things).* Methods in Ecology and Evolution, 2012. **3**(2): p. 217-223.

26. Garland, T., Jr., et al., *Phylogenetic Analysis of Covariance by Computer Simulation.* Systematic Biology, 1993. **42**(3): p. 265-292.

27. Khabbazian, M., et al., *Fast and accurate detection of evolutionary shifts in Ornstein–Uhlenbeck models.* Methods in Ecology and Evolution, 2016. **7**(7): p. 811-824.

28. Health Care Productivity, I. and L. Minitab, *RANDOM FORESTS*. 2006.

29. Tin Kam, H. *Random decision forests*. in *Proceedings of 3rd International Conference on Document Analysis and Recognition*. 1995.

30. Fürnkranz, J., *Decision Tree*, in *Encyclopedia of Machine Learning*, C. Sammut and G.I. Webb, Editors. 2010, Springer US: Boston, MA. p. 263-267.

31. Lucas, T.C.D., *A translucent box: interpretable machine learning in ecology.* Ecological Monographs, 2020. **90**(4): p. e01422.

32. Benito, M., *spatialRF: Easy spatial regression with random forest.* R package version 1.1.0, 2021. **1**(0).

33. Kuhn, M. and H. Wickham, *Tidymodels: a collection of packages for modeling and machine learning using tidyverse principles.* 2020.

34. Haibo, H., et al. *ADASYN: Adaptive synthetic sampling approach for imbalanced learning.* in *2008 IEEE International Joint Conference on Neural Networks (IEEE World Congress on Computational Intelligence)*. 2008.

35. Liaw, A. and M. Wiener, *Classification and regression by randomForest.* R news, 2002. **2**(3): p. 18-22.

36. Hand, D.J. and R.J. Till, *A Simple Generalisation of the Area Under the ROC Curve for Multiple Class Classification Problems.* Machine Learning, 2001. **45**(2): p. 171-186.

37. Molnar, C., *Interpretable Machine Learning: A guide for Making Black Box Models Explanable (2nd ed.).* 2022, christophm.github.io/interpretable-ml-book/.

38. Greenwell, B.M., B.C. Boehmke, and B. Gray, *Variable Importance Plots-An Introduction to the vip Package.* R J., 2020. **12**(1): p. 343.

39. Uyeda, J.C., D.S. Caetano, and M.W. Pennell, *Comparative Analysis of Principal Components Can be Misleading.* Systematic Biology, 2015. **64**(4): p. 677-689.

40. Ducatez, S. and D.J. Field, *Disentangling the avian altricial-precocial spectrum: Quantitative assessment of developmental mode, phylogenetic signal, and dimensionality.* Evolution, 2021. **75**(11): p. 2717-2735.

41. Beaulieu, J.M., et al., *MODELING STABILIZING SELECTION: EXPANDING THE ORNSTEIN–UHLENBECK MODEL OF ADAPTIVE EVOLUTION.* Evolution, 2012. **66**(8): p. 2369-2383.
